# IRE1 drives a homeostatic response to reduced protein influx into the endoplasmic reticulum

**DOI:** 10.64898/2026.03.02.709157

**Authors:** Francesca Zappa, Advait Subramanian, Benjamin Yang, Julia E. Conrad, Tsan-Wen Lu, Jing Wang, Nicolas A. DeBeaubien, Rose Yan, Renato Minopoli, Tristan Croll, Stefka Tyanova, Mauro Costa-Mattioli, Peter Walter, Diego Acosta-Alvear

**Author notes:** These authors contributed equally (Francesca Zappa, Advait Subramanian). Department of Biochemistry and Biophysics, University of California at San Francisco, San Francisco, CA, USA.

## Abstract

IRE1, alongside ATF6 and PERK, orchestrates the Unfolded Protein Response, a network of signaling pathways that maintains endoplasmic reticulum (ER) homeostasis. Two modes of IRE1 activation are known: i) in response to an accumulation of unfolded proteins in the ER lumen and ii) in response to compositional changes to the ER membrane that alter its physical properties. Here we identify a third, independent mode of IRE1 activation: ER co-translational translocation deficits activate IRE1 through a mechanism that relies on the release of IRE1 molecules from unoccupied translocons. We define this mechanism as TRES for “TRanslocon Engagement Surveillance”. TRES leads to spontaneous activation of IRE1 and bypasses its unfolded protein- and ER membrane composition-sensing functions. Inhibiting translation initiation similarly activates IRE1 by TRES, as it leads to a decline in ER protein import, thus linking the Integrated Stress Response to IRE1 signaling. TRES drives IRE1 activation without activating ATF6 or PERK, resulting in a distinct gene expression program that feeds back by boosting the co-translational translocation machinery to rebalance the ER protein load. Our findings thus demonstrate that monitoring and adjusting the rates of protein translocation are critical for maintaining ER homeostasis.

## Introduction

Surveillance of protein targeting and import across organelle membranes is a fundamental process that ascertains that the protein folding load and the available protein folding machinery are properly balanced. In the endoplasmic reticulum (ER), protein targeting is facilitated by the signal recognition particle (SRP), which delivers translating ribosomes synthesizing signal sequence-bearing peptides to the translocon, initiating the co-translational translocation of nascent polypeptide chains into the ER lumen (*1–3*). The translocon is a multiprotein complex composed of SEC61α, β, and γ subunits and accessory factors (*4*). Dedicated ER quality control mechanisms counterbalance deviations from homeostasis to maintain proper ER physiology and adjust the organelle’s functions according to need. A key ER quality control mechanism is the Unfolded Protein Response (UPR). The UPR is a signaling network that detects protein folding defects (“unfolded protein stress”) as well as changes in ER membrane composition and fluidity (“lipid bilayer stress”) to activate transcriptional programs that rebalance ER function (*5–7*). In animals, the UPR is governed by three ER membrane-resident sensors: the kinase PERK, the transcription factor ATF6, and the kinase/RNase IRE1. Of the three sensors, only IRE1 interacts with the SRP, the ribosome, and the translocon (*8*, *9*), yet the functional implications of these associations remain largely unclear (*8–15*).

Current models posit that IRE1 activation results from the direct binding of unfolded protein ligands to its ER lumenal domain (*16–19*), the reversible dissociation of the lumenal ER chaperone BiP/GRP78 from IRE1 (*20*), and the sensing of lipid bilayer stress by IRE1’s transmembrane domain (*21–23*). In response to these perturbations, IRE1 molecules self-associate and *trans*-autophosphorylate, both a prerequisite for awakening its endoribonuclease activity. Active IRE1 initiates the non-conventional splicing of the *X-box binding protein 1* (*XBP1*) mRNA to generate the central UPR transcription factor XBP1s (“s” for spliced) (*24*) that upregulates the transcription of genes involved in protein folding, protein export, ER-associated degradation (ERAD), and lipid biosynthesis (*25–29*) in order to rebalance the ER’s functional capacity. In addition, activated IRE1 degrades ER-localized mRNAs through RIDD (regulated IRE1-dependent decay) (*30*) and RIDDLE (RIDD <u>L</u>acking <u>E</u>ndomotif) (*31*), thereby reducing the flux of proteins entering the ER. Through these actions, IRE1 mitigates ER stress (*32–34*). If this cannot be achieved and ER stress remains unmitigated, sustained IRE1 activity induces cell death (*35–37*). Given its fundamental role in making life/death decisions, it comes as no surprise that maladaptive IRE1 signaling is linked to the pathophysiology of multiple neurodegenerative, metabolic, and malignant diseases (*7*).

Along with RIDD and RIDDLE, the flux of proteins entering the ER lumen can also be adjusted by controlling protein synthesis. A prominent mechanism governing protein synthesis rates is the Integrated Stress Response (ISR). The ISR is a homeostatic signaling network that detects a myriad of stresses, including unfolded protein stress, and temporarily curtails protein synthesis by attenuating translation initiation (*38*). Translational control via the ISR can be imposed on a rapid time scale and thus provides the cell with enough time to adapt through changes in gene expression. The ISR is thus intimately intertwined with the UPR, as both stress responses share a common sensor, PERK. Besides PERK, three additional ISR stress-sensing kinases, GCN2, HRI and PKR, converge on phosphorylating the alpha subunit of eukaryotic initiation factor 2 (eIF2α). Phosphorylation of eIF2α leads to the translational suppression of the vast majority of mRNAs and consequently limits the protein load inside the ER (*38*, *39*).

Here, we describe TRES (for <u>TR</u>anslocon <u>E</u>ngagement <u>S</u>urveillance) as a third and heretofore unrecognized mode of IRE1 activation that operates independent of IRE1’s unfolded protein and lipid bilayer stress sensing modalities. TRES couples IRE1 activation to decreased ER protein influx. IRE1 activation by TRES results from co-translational translocation deficits elicited by insufficient ER protein targeting, translocon dysfunction, and fluctuating translation initiation rates, as occur upon induction of the ISR. These deficits dissociate IRE1 from the translocon, allowing for its activation. The transcriptional output resulting from TRES-activated IRE1 signaling is distinct from that of the classically defined UPR. The discovery of TRES thus expands the concept of ER stress to include conditions resulting in ER *under*loading. These findings establish that disrupted protein translocation into the ER is a distress signal sensed by IRE1.

## Results

### Dissociation from the translocon derepresses IRE1 during TRES

To elucidate the functional interplay between IRE1α (IRE1 hereafter) and the ER co-translational translocation machinery, we generated four mNeonGreen-tagged and doxycycline (Dox)-inducible expression constructs (Fig. 1A):

i) IRE^WT^ (wild-type IRE1);
ii) IRE1^TAD^ (for “translocon association deficient”). IRE1 with a ten amino acid deletion, ΔE434-D443, that reduces its association with the translocon (*10*) (Fig. S1A);
iii) IRE1^DAB^ (for “deaf and blind”). IRE1 with a core lumenal domain deletion, ΔI19-D408, which blocks its ability to bind to unfolded proteins and BiP (“blind”), combined with a point mutation, W457A, that renders it unable to sense lipid bilayer stress (“deaf”) (*22*, *23*) (Fig. S1B); and
iv) IRE1^BTAD^ (for “blind and translocon association deficient”). IRE1 with a large deletion that removes its entire lumenal portion, including its translocon-binding motif, ΔI19-D443, which prevents unfolded protein, BiP, and translocon binding but retains lipid bilayer stress sensing (Fig. S1C).

**Figure 1.**
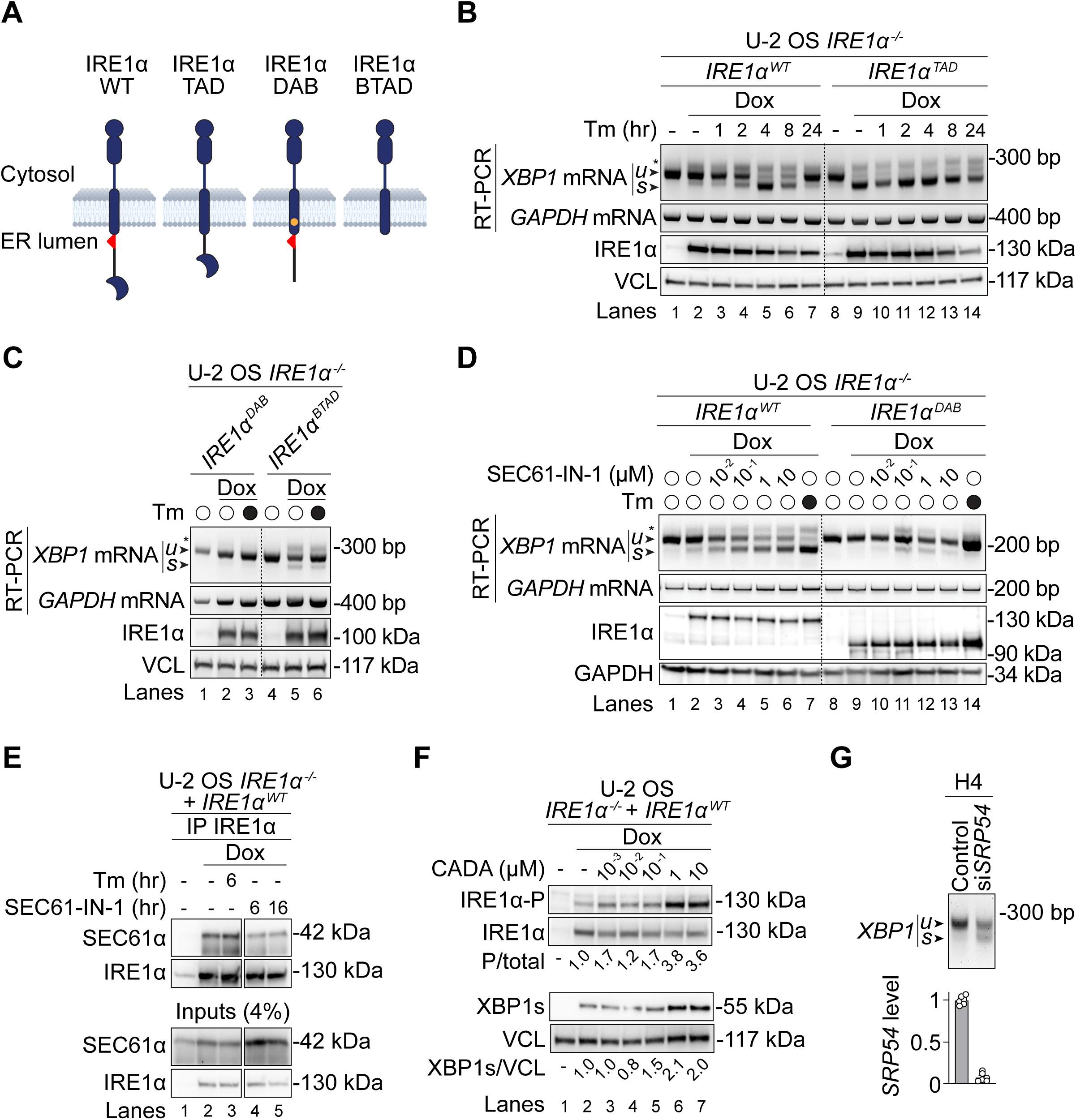
Disrupted ER translocation activates IRE1. (A) Schematic representation of the different IRE1 constructs used in this study. Solid red triangle (a.a. E434-D443): translocon association motif; orange dot: W457A. Each doxycycline-inducible transgene was inserted as a single copy using the Flp-In system in U-2 OS *IRE1α^-/-^* cells. (B) RT-PCR of *XBP1* and *GAPDH* mRNAs (top), immunoblotting of IRE1α, and vinculin (VCL) (bottom) in lysates obtained from U-2 OS *IRE1α*^-/-^ cells expressing mNeonGreen-tagged IRE1α^WT^ and IRE1α^TAD^ upon addition of 5 ng/mL doxycycline (Dox). Cells were treated with 5 µg/mL tunicamycin (Tm) for the indicated times. u - unspliced; s - spliced *XBP1* mRNA amplicons, respectively. The asterisk indicates a hybrid amplicon generated by annealing of one strand of *XBP1u* and one strand of *XBP1s* during the PCR. *GAPDH*, VCL: loading controls. Quantification of the percentage of *XBP1* mRNA splicing is shown in Fig. S1F. Data are representative of three independent experiments. (C) RT-PCR and immunoblots in lysates obtained from U-2 OS *IRE1α*^-/-^ cells expressing mNeonGreen-tagged IRE1α^DAB^ and IRE1α^BTAD^ upon addition of 5 ng/mL Dox. Cells were treated with 5 µg/mL Tm for 4 hours where indicated. Quantification of the percentage of *XBP1* mRNA splicing is shown in Fig. S1G. Data are representative of three independent experiments. (D) RT-PCR and immunoblots in lysates obtained from U-2 OS *IRE1α*^-/-^ cells expressing mNeonGreen-tagged IRE1α^WT^ and IRE1α^DAB^ upon addition of 5 ng/mL doxycycline (Dox). Cells were treated with SEC61-IN-1 for 16 hours at the indicated concentrations. Quantification of the percentage of *XBP1* mRNA splicing is shown in Fig. S2A. Data are representative of three independent experiments. (E) Immunoprecipitation of mNeonGreen-tagged IRE1α^WT^ in U-2 OS *IRE1α*^-/-^ cells expressing IRE1α^WT^ upon addition of 5 ng/mL Dox. Cells were treated with 5 µg/mL Tm for 6 hours or 1 µM SEC61-IN-1 for 6 and 16 hours as indicated. Immunoblotting was performed using antibodies against SEC61α and IRE1α. Lanes are cropped from images of the same gel. Data are representative of two independent experiments. (F) Immunoblots of phosphorylated and total IRE1α (top) and spliced XBP1 protein and vinculin (bottom) in lysates obtained from U-2 OS *IRE1α*^-/-^ cells expressing mNeonGreen-tagged IRE1α^WT^ upon addition of 5 ng/mL Dox. Cells were treated with CADA for 16 hours at the indicated concentrations. The quantification of the proportion of phosphorylated IRE1 and spliced XBP1 are shown below the immunoblots. Data are representative of two independent experiments. VCL: loading control. (G) RT-PCR of *XBP1* mRNA in H4 cells depleted of *SRP54* by RNA interference. Quantification of the percentage of *XBP1* mRNA splicing is shown in Fig. S2D. Data are representative of three independent experiments. Bottom panel: extent of *SRP54* knockdown measured by RT-qPCR and normalized internally to *GAPDH*. Mean ± SEM expressed as fold change over the levels in mock-transfected controls (N=3).

The constructs were integrated into *IRE1^-/-^* U-2 OS (osteosarcoma) cells at a defined genomic locus using the Flp-In system (*40*), thereby allowing comparable and regulated transgene expression. Dox treatment (5 ng/mL) induced robust expression of the transgenes, which we estimated to be between ∼8- and ∼16-fold over endogenous IRE1 protein levels observed in parental U-2 OS cells (Fig. S1D-E) (*41*).

We monitored the dynamics of IRE1 activation in *IRE1^-/-^*cells expressing IRE1^WT^ or IRE1^TAD^ by challenging them with the *N*-linked glycosylation inhibitor tunicamycin (Tm), an ER stress-inducing agent that promotes the buildup of unfolded proteins in the ER lumen. Expression of IRE1^WT^ led to a basal *XBP1* mRNA splicing (26±9%) (Fig. 1B; lane 2, Fig. S1F). This signal, observed in the absence of ER stress, is a result of the overexpression level of the transgene. As expected from previous observations, *XBP1* mRNA splicing was boosted after Tm treatment (82±8% at 4 hours) (Fig. 1B; lane 5, Fig. S1F), before attenuating back to baseline (28±14% at 24 hours) (Fig. 1B; lane 7, Fig. S1F) (*42*). Unexpectedly, expression of IRE1^TAD^ in the absence of ER stress led to high *XBP1* mRNA splicing (77±4%) (Fig. 1B, lane 9, Fig. S1F). By contrast to the *XBP1* mRNA splicing fold-changes (∼4-fold) observed for IRE1^WT^ (Fig. 1B, compare lanes 2 and 5, Fig. S1F), ER stress induction in cells expressing IRE1^TAD^ did not induce substantive *XBP1* mRNA splicing (∼1.1-fold over wild-type) (Fig. 1B; compare lanes 9 and 12, Fig. S1F). Moreover, IRE1^TAD^ activity remained elevated and, surprisingly, did not attenuate as observed for IRE1^WT^ (Fig. 1B; compare lanes 9 and 14, Fig. S1F). Both IRE1^WT^ and IRE1^TAD^ abundance gradually decreased to similar degrees over the time course of the experiment. Thus, a decrease in IRE1 protein levels does not explain the differences in activity observed between the wild type and TAD mutant.

To assess the contributions of IRE1’s unfolded protein and ER membrane stress sensing, we next expressed IRE1^BTAD^ and IRE1^DAB^ and measured *XBP1* mRNA splicing. The expression of IRE1^BTAD^, which cannot bind to the translocon, was necessary and sufficient to induce *XBP1* mRNA splicing both in the absence and after induction of ER stress. However, splicing was less prominent than shown above for IRE1^TAD^ (28±2%) (Fig. 1C, lanes 4-6, Fig. S1G), consistent with the notion that the lumenal domain potentiates IRE1 multimerization, which results in signal amplification (*18*, *19*). By contrast, the expression of IRE1^DAB^, which can bind to the translocon, failed to induce *XBP1* mRNA splicing both in the absence and after induction of ER stress (Fig. 1C; lanes 1 – 3, Fig. S1G).

Together, the data presented so far suggest that IRE1 is derepressed upon dissociation from the translocon (IRE1^TAD^ and IRE1^BTAD^) and that the activity resulting from this brake-release is independent of IRE1’s lumenal unfolded protein sensing domain.

### IRE1 senses co-translational translocation defects during TRES

Previous work indicated that depleting SEC61 subunits or clogging translocons with artificial reporter constructs activate IRE1 (*12*, *15*). Accordingly, translocon inhibitors induce IRE1 activation in cells (*43–46*). To address whether a reduction of co-translational translocation into the ER results in IRE1 activation by way of its dissociation from the translocon, we disrupted ER co-translational translocation and measured *XBP1* mRNA splicing resulting from IRE1 activation.

First, we pharmacologically inhibited the translocon with the small-molecule inhibitor SEC61-IN-1 (compound A317 in (*47*)) in cells expressing IRE1^WT^. These experiments showed a dose-dependent increase in *XBP1* mRNA splicing (∼2-fold to ∼3.5-fold over baseline) (Fig. 1D; lanes 3-6, Fig. S2A) with an accompanying reduction in the association between IRE1 and SEC61𝛼 (Fig. 1E; compare lanes 2, 4, and 5). By contrast, Tm treatment induced *XBP1* mRNA splicing without affecting the interaction between IRE1 and SEC61𝛼, as previously described (*9*, *48*) (Fig. 1E; lanes 2-3). Strikingly, SEC61-IN-1 treatment also induced *XBP1* mRNA splicing in *IRE1^-/-^* cells expressing IRE1^DAB^ (25±3%) (Fig. 1D, lanes 11-12, Fig. S2A), similar to that observed upon expression of IRE1^BTAD^ (28±2%) (compare Fig. 1C, lanes 5 and 6, with Fig. 1D lanes 11 and 12). Taken together, these data indicate that neither unfolded-protein binding, nor BiP-dissociation, nor lipid bilayer stress sensing is required for IRE1 activity upon treatment with SEC61-IN-1.

To confirm these conclusions, we treated cells with a different translocon inhibitor, cyclotriazadisulfonamide (CADA), for which the molecular mechanism of action has been defined structurally (*49*). Molecular docking onto SEC61𝛼 shows that CADA occupies a pocket nested between the plug and the lateral gate of the channel (Fig. S2B, top), as does SEC61-IN-1 (Fig. S2B, bottom). These analyses suggest that both drugs occupy a similar pocket and stabilize the translocon in a conformation incompatible with translocation. Indeed, CADA activated IRE1^WT^, as shown by IRE1 autophosphorylation and the accumulation of XBP1s protein (Fig. 1F). These observations were not restricted to our transgene experiments, as SEC61-IN-1 and CADA induced XBP1s protein accumulation in parental U-2 OS cells expressing endogenous IRE1 (Fig. S2C).

IRE1 activation involves the formation of higher order oligomers (*42*). Whereas Tm treatment expectedly induced IRE1 foci (Fig. S3, arrows) in U-2 OS cells expressing fluorescently tagged IRE1^WT^, treatment with SEC61-IN-1 did not (Fig. S3). These results suggest that large IRE1 assemblies do not form during TRES.

To orthogonally test the notion that inhibiting translocon activity induces IRE1, we disrupted co-translational translocation by depleting the SRP subunit SRP54 in a different cell line solely relying on endogenous levels of IRE1. We reasoned that preventing ribosome targeting to the translocon in cells depleted of SRP would lead to IRE1 activation. Indeed, SRP54 depletion with small interfering RNAs in H4 neuroglioma cells resulted in *XBP1* mRNA splicing (32±8%) (Fig. 1G, S2D), corroborating previous results (*12*).

In a third orthogonal approach, we reduced ER translocation by interrupting general translation initiation which impedes co-translational ribosome targeting to the ER. To this end, we treated cells with the translation initiation inhibitors rocaglamide A (RocA) and silvestrol, which clamp the eIF4A RNA helicase to 5’ untranslated regions of mRNAs and prevent efficient 43S ribosomal subunit scanning (*50*, *51*). Treatment with RocA resulted in a reduced ribosome density on ER membranes, as confirmed by electron microscopy (Fig. S4A). RocA treatment led to the dissociation of IRE1^WT^ from the translocon and from ribosomal proteins (Fig. 2A), with a corresponding induction of *XBP1* mRNA splicing (95±5%) (Fig. 2B, lane 5, Fig. S4B). RocA treatment similarly led to the dissociation of IRE1^DAB^ from the translocon and ribosomal proteins (Fig. 2C) and IRE1^DAB^ dependent *XBP1* mRNA splicing (45±5%, lane 5) (Fig. 2D, Fig. S4C).

**Figure 2.**
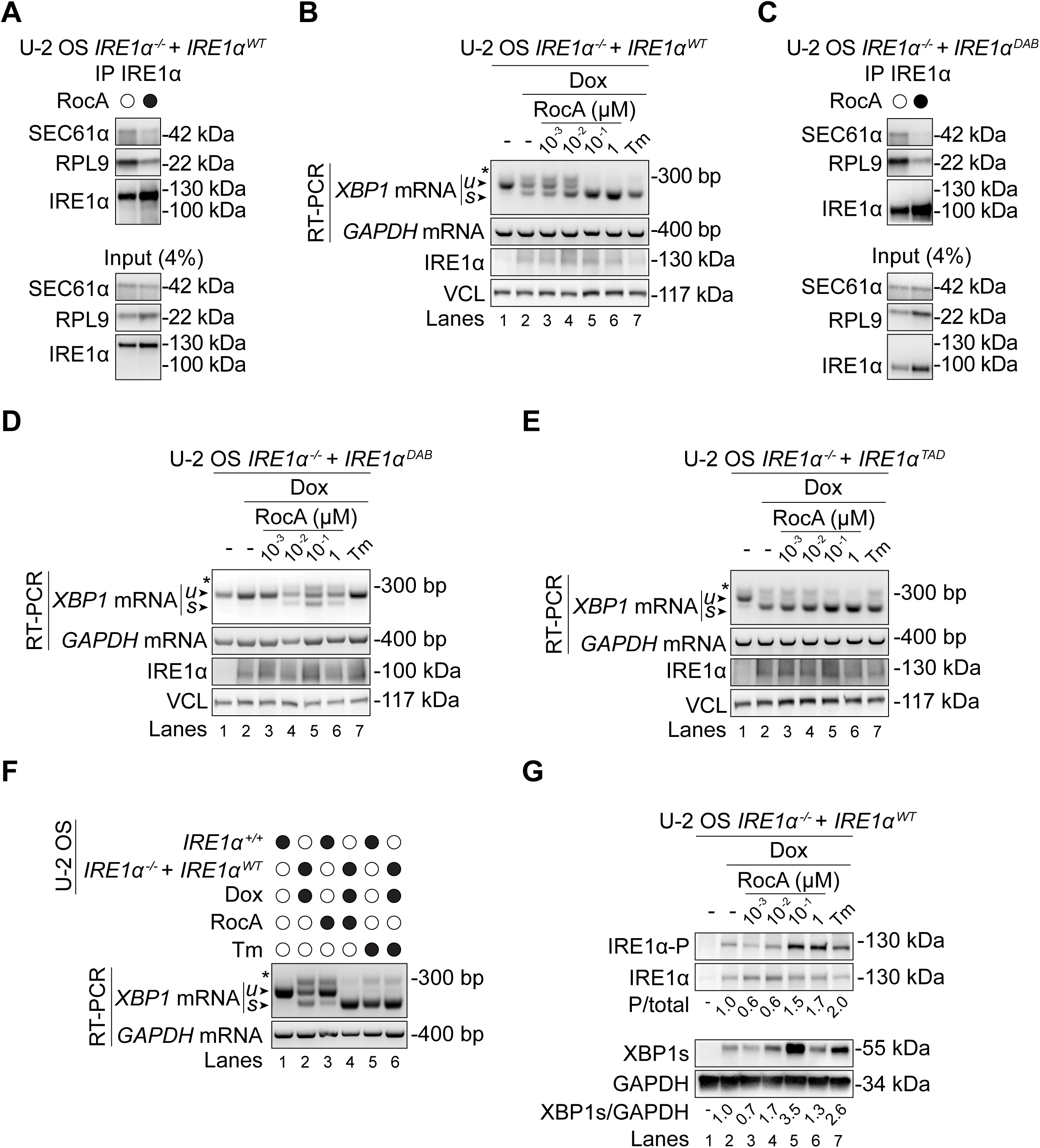
Translation initiation inhibition activates IRE1. (A) Immunoprecipitation of mNeonGreen-tagged IRE1α^WT^ in U-2 OS *IRE1*α^-/-^ cells expressing IRE1α^WT^ upon addition of 5 ng/mL Dox. Cells were treated with 500 nM RocA for 16 hours as indicated. Immunoblotting was performed using antibodies against SEC61α, RPL9, and IRE1α. Data are representative of two independent experiments. (B) RT-PCR of *XBP1* and *GAPDH* mRNAs (top), immunoblotting of IRE1α, and VCL (bottom) in lysates obtained from U-2 OS *IRE1α*^-/-^ cells expressing mNeonGreen-tagged IRE1α^WT^ upon addition of 5 ng/mL Dox. Cells were treated with RocA for 16 hours at the indicated concentrations or 5 µg/mL Tm for 4 hours. u - unspliced; s - spliced *XBP1* mRNA amplicons, respectively. The asterisk indicates a hybrid amplicon generated by annealing of one strand of *XBP1u* and one strand of *XBP1s* during the PCR. *GAPDH*, VCL: loading controls. Quantification of the percentage of *XBP1* mRNA splicing is shown in Fig. S4B. Data are representative of three independent experiments. (C) Immunoprecipitation of mNeonGreen-tagged IRE1α^DAB^ as in panel A. (D-E) RT-PCR and immunoblotting as in panel B of lysates obtained from U-2 OS *IRE1α*^-/-^ cells expressing mNeonGreen-tagged IRE1α^DAB^ (D), and mNeonGreen-tagged IRE1α^TAD^ (E). Quantification of the percentage of *XBP1* mRNA splicing is shown in Figs. S4C-D. Data are representative of three independent experiments. (F) RT-PCR of *XBP1* and *GAPDH* mRNAs in lysates obtained from U-2 OS *IRE1α*^+/+^ cells and U-2 OS *IRE1α*^-/-^ cells expressing IRE1α^WT^ cells upon addition of 5 ng/mL Dox. Cells were treated with 100 nM RocA for 16 hours or 5 µg/mL Tm for 4 hours as indicated. Data are representative of three independent experiments. (G) Immunoblots of phosphorylated and total IRE1α (top) and spliced XBP1 protein and VCL (bottom) in lysates obtained from U-2 OS *IRE1α*^-/-^ cells expressing mNeonGreen-tagged IRE1α^WT^ upon addition of 5 ng/mL Dox. Cells were treated with RocA for 16 hours at the indicated concentrations or 5 µg/mL Tm for 4 hours as indicated. The quantification of the proportion of phosphorylated IRE1 and spliced XBP1 are shown below the immunoblots. Data are representative of three independent experiments. GAPDH: loading control.

Notably, RocA treatment minimally affected the already high *XBP1* mRNA splicing activity of the translocon-binding-deficient IRE1^TAD^ mutant (Fig. 2E, Fig. S4D), indicating that IRE1 dissociation from the translocon is sufficient for high IRE1 activity. Accordingly, RocA treatment increased *XBP1* mRNA splicing by IRE1^WT^ to a degree comparable to that elicited by Tm treatment (Fig. 2F, compare lanes 4-6), suggesting that disrupting co-translational translocation activates IRE1^WT^ to the same extent as ER stress. Indeed, RocA treatment induced IRE1 autophosphorylation (Fig. 2G) and XBP1s protein accumulation (Fig. 2G, S4E), and RIDD (Fig. S4F). As we observed upon SEC61-IN-1 treatment, RocA did not induce prominent IRE1 foci (Fig. S4H). High RocA concentrations (1 µM) did not elevate XBP1s protein levels despite inducing *XBP1* mRNA splicing due to a strong translation initiation block (Fig. S4G).

Similar to our experiments with SEC61-IN-1 and CADA, RocA treatment of U-2 OS cells expressing endogenous IRE1 elicited *XBP1* mRNA splicing, albeit modestly when compared to IRE1^WT^ overexpression (Fig. 2F, compare lanes 1 and 3, and lanes 2 and 4). Treatment with Tm, however, induced high *XBP1* mRNA splicing in these cells (Fig. 2F, compare lanes 1 to 2 and 5 to 6). We obtained similar results in H4 neuroglioma cells treated with silvestrol (Fig. S5A) and in *ex vivo* liver tissues of mice treated with Zotatifin, a synthetic RocA analogue (Fig. S5B). Notably, and by contrast to our observations with RocA and silvestrol, we detected no *XBP1* mRNA splicing after treating U-2 OS cells with the elongation inhibitors cycloheximide (CHX) and emetine (EME), which would stabilize ribosome-nascent chain complexes on translocons (*52*) (Fig. S5C).

Notably, neither ATF6 nor PERK were activated by RocA, SEC61-IN-1 or CADA (Fig. S5D), which as expected were both activated by the classical UPR inducer dithiothreitol (DTT), a reducing agent that disrupts disulfide bonds, as indicated by ATF6 proteolytic processing and a change in PERK’s electrophoretic mobility in SDS-PAGE due to its autophosphorylation (Fig. S5D). We noted that the highest concentration of SEC61-IN-1 reduced the protein levels of GFP-ATF6 and PERK (Fig. S5D, lane 12), but not IRE1 (Fig. S2C, lane 5). It is possible that PERK and ATF6 molecules are no longer inserted into the ER membrane under these conditions (*53*) or that they are selectively degraded.

Taken together, our genetic and pharmacological experiments converge on a model in which co-translational translocation defects preferentially activate IRE1 by promoting its dissociation from SEC61. This activation modality is independent of IRE1’s canonical unfolded protein binding or lipid bilayer stress-sensing.

### The Integrated Stress Response selectively activates IRE1 by TRES

The findings above demonstrate that reduced translation initiation activates IRE1. These observations raise the intriguing possibility that the ISR, which operates as a master regulator of translation initiation and intimately connects to the UPR, could activate IRE1. To address this notion, we activated the ISR directly using FKBP-PKR, an engineered ISR kinase that can be activated by the drug-like small molecule FKBP homodimerizer AP-20187 and leads to suppression of translation initiation (Fig. 3A, S6A) (*54*, *55*). We chose PKR because it responds to viral infection and not to unfolded protein buildup.

**Figure 3.**
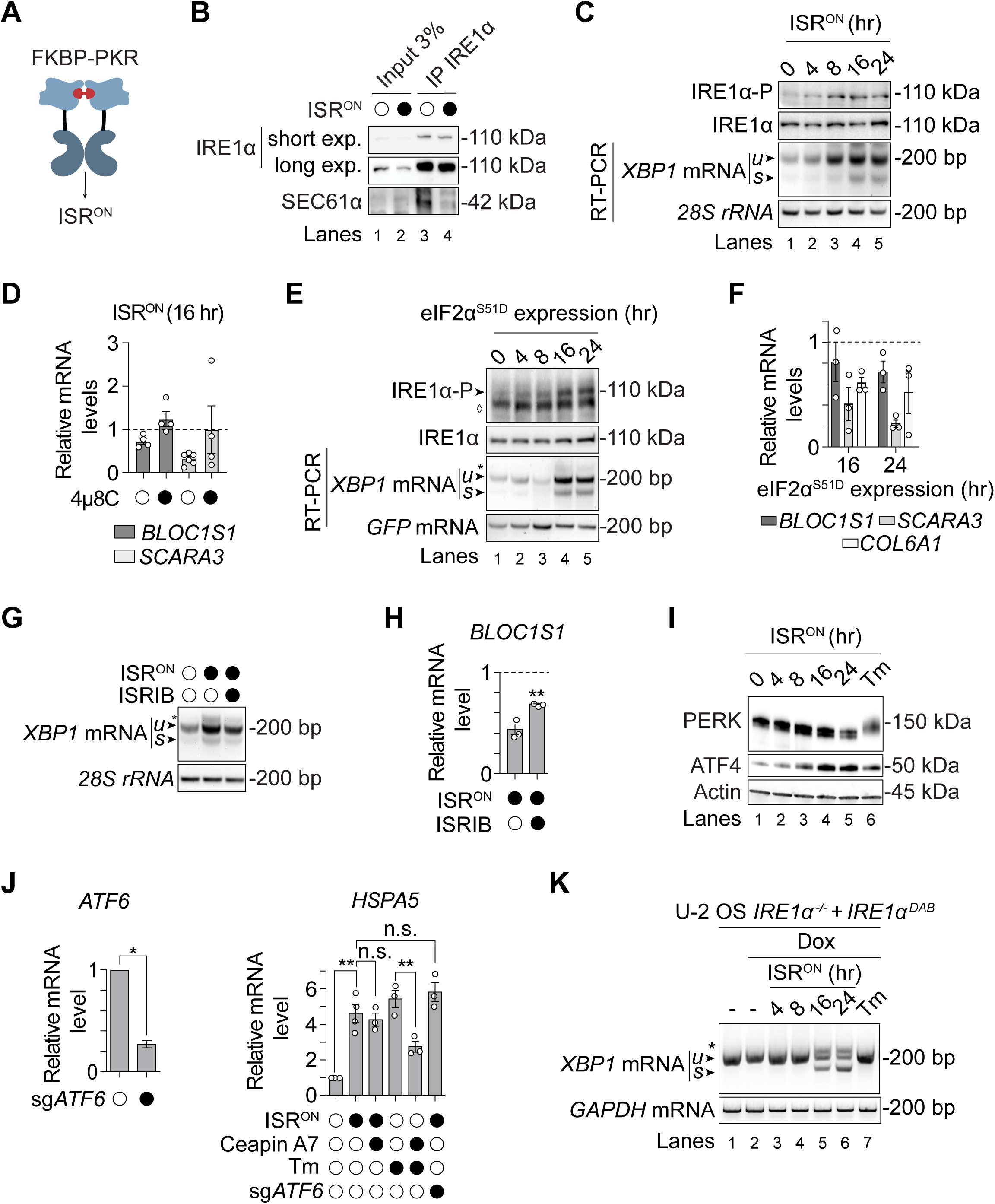
Synthetic activation of the ISR induces IRE1. (A) Schematic representation of FKBP-PKR. The homodimerizer is shown in red. The FKBP^F36V^ domain (light blue) is adjoined to the PKR kinase domain (dark blue) by PKR’s unstructured linker. (B) Co-immunoprecipitation of IRE1α and SEC61α in lysates obtained from H4 cells expressing FKBP-PKR and treated with the homodimerizer for 16 hours. (C) Immunoblot of IRE1α phosphorylation (IRE1α-P) and RT-PCR of *XBP1* mRNA and *28S* ribosomal RNA in lysates obtained from H4 cells expressing FKBP-PKR and treated with the homodimerizer for the indicated times. Loading control: *28S* ribosomal RNA. Data are representative of five independent experiments. (D) RT-qPCR of *BLOC1S1* and *SCARA3* mRNAs in H4 cells expressing FKBP-PKR and treated with the homodimerizer for 16 hours with or without co-treatment with 10 µM of the IRE1 RNase inhibitor 4µ8C. Data: Mean ± SEM of fold changes normalized to the levels in untreated controls (N≥3). (E) Immunoblot of IRE1α phosphorylation (IRE1α-P) and RT-PCR of *XBP1* and *GFP* mRNAs in lysates obtained from H4 cells expressing eIF2α^S51D^ hosted in a construct that also expresses GFP, for the indicated times. *GFP* mRNA: transfection and loading control. Data are representative of three independent experiments. The ♢ indicates a non-specific band in the immunoblot (top) and * denotes the hybrid *XBP1* amplicon in the RT-PCR (bottom). (F) RT-qPCR of *BLOC1S1* and *SCARA3* mRNAs in H4 cells expressing eIF2α^S51D^ for 16 hours. Data: Mean ± SEM of fold changes normalized to the levels in untreated controls (N = 3). (G) RT-PCR of *XBP1* mRNA and *28S* ribosomal RNA in H4 cells expressing FKBP-PKR and treated with the homodimerizer for 16 hours with or without co-treatment with 800 nM ISRIB. Data are representative of three independent experiments. (H) RT-qPCR of the expression levels of the *BLOC1S1* mRNA in H4 cells expressing FKBP-PKR, treated with the homodimerizer for 16 hours with or without addition of 800 nM ISRIB. Data: Mean ± SEM of fold changes normalized to the respective levels in untreated controls (N = 3). **p-value < 0.01, Student’s t-test. (I) Immunoblots of PERK and ATF4 in lysates obtained from H4 cells expressing FKBP-PKR and treated with the homodimerizer for the indicated times and 5 µg/ml Tm for 4 hours. Actin: loading control. (J) left panel: RT-qPCR of the *ATF6* mRNA in H4 cells with CRISPRi-mediated knockdown of *ATF6* (sgATF6); right panel: *HSPA5* mRNA in H4 cells expressing FKBP-PKR cells under the following conditions: 2.5 µg/ml Tm for 16 hours, 5 nM Ceapin-A7, and homodimerizer treatment for 16 hours, as indicated, or in cells with CRISPRi-mediated knockdown of ATF6 (sgATF6). For the right panel data: Mean ± SEM of fold changes normalized to the levels in untreated controls. **p-value <0.01, Student’s t-test; n.s. - not significant. (K) RT-PCR of *XBP1* and *GAPDH* mRNAs in U-2 OS *IRE1α*^-/-^ cells expressing FKBP-PKR and mNeonGreen-tagged IRE1α^DAB^ treated with 20 ng/mL Dox for 48 hours, 5 µg/mL Tm for 4 hours as indicated, and the homodimerizer for the indicated times. Data are representative of three independent experiments. The asterisk indicates the hybrid *XBP1* amplicon in the RT-PCR.

Indeed, FKBP-PKR activation with AP-20187 (referred to as ISR^ON^ hereafter) activated IRE1 as assessed by multiple lines of evidence:

i) it disrupted the IRE1-translocon interaction (Fig. 3B, compare lanes 3 and 4);
ii) it induced IRE1 autophosphorylation (Fig. 3C – lanes 4 and 5);
iii) it induced *XBP1* mRNA splicing (Fig. 3C – lanes 4 and 5; S6B – lanes 5 and 6) and upregulated XBP1s transcriptional target genes (Fig. S6C,D); and
iv) it activated RIDD, which was suppressed upon co-treatment with the IRE1 RNase inhibitor 4µ8C (Fig. 3D).

Because the ISR signals through eIF2 phosphorylation (*38*), and to rule out potential off-target effects of FKBP-PKR activation, we next overexpressed a phosphomimetic mutant of the α-subunit of eIF2 bearing the S51D mutation, eIF2α^S51D^ (*54*). As occurred with our ISR^ON^ approach, overexpression of eIF2α^S51D^ (Fig. S6E) was sufficient to induce IRE1 autophosphorylation and *XBP1* mRNA splicing (Fig. 3E), upregulate XBP1s transcriptional targets (Fig. S6F, G), and activate RIDD (Fig. 3F), corroborating that the ISR activates IRE1. Accordingly, treatment with ISRIB, a small molecule that renders cells refractory to the effects of eIF2 phosphorylation (*56*), reduced *XBP1* mRNA splicing (Fig. 3G) and XBP1s target expression (Fig. S6H, I), and diminished RIDD (Fig. 3H) in ISR^ON^ cells.

As a third approach to induce the ISR, we induced mitochondrial stress to activate the ISR kinase HRI with mitochondrial uncouplers (oligomycin and BTdCPU) (*57–59*). We observed induction of *XBP1* mRNA splicing that was reduced by treatment with ISRIB and 4µ8C (Fig. S6J, K). We note that ISR activation increased total *XBP1* mRNA levels, in addition to activating *XBP1* mRNA splicing (Fig. 3C, 3E); both these effects were dampened by ISRIB treatment (Fig. 3G). These results indicate that *XBP1* mRNA abundance is autoregulated by the ISR (*25*), in addition to being regulated by ATF6 upon UPR induction (*24*).

To rule out potential involvement of the other UPR sensors under ISR^ON^ conditions, we measured the electrophoretic mobility of PERK and the induction of ATF6’s canonical target genes (*HSPA5/BiP* and *GRP94*). We observed no changes in PERK’s electrophoretic mobility in SDS-PAGE (Fig. 3I), suggesting that PERK does not become phosphorylated. Similarly, the level of the *GRP94* mRNA remained unchanged (Fig. S6L). By contrast, ISR^ON^ elevated *HSPA5* mRNA levels (Fig. 3J and S6C). This result is easily reconciled because *HSPA5* transcription is regulated by both ATF6 and XBP1s (*25*, *26*, *60*). To demonstrate directly the lack of ATF6’s involvement in ISR^ON^, we depleted ATF6 using CRISPRi or alternatively blocked its activation with Ceapin-A7, a small molecule ATF6 inhibitor (Fig. 3J) (*61*). Neither manipulation affected the *HSPA5* mRNA levels upon ISR^ON^ induction (Fig. 3J, right panel - compare bar 2 to bars 3 and 6). As expected for the effect solely resulting from ISR activation, ISRIB treatment blunted *HSPA5* mRNA upregulation upon ISR^ON^ (Fig. S6I) confirming that ISR^ON^ does not activate PERK or ATF6. Further supporting these conclusions, ISR^ON^ led to *XBP1* mRNA splicing by IRE1^DAB^ (Fig. 3K) and did not induce IRE1^WT^ foci (Fig. S6M).

Together, these data indicate that the translation initiation block and consequential co-translational protein translocation deficits imposed by the ISR lead to a selective activation of IRE1 that do not require IRE1’s canonical ER stress-sensing modalities or the formation of large IRE1 oligomers.

### The IRE1-induced gene expression program differs between TRES and the UPR

The selective activation of IRE1 in response to co-translational translocation deficits prompted us to ask whether IRE1 induces customized gene expression programs to match specific physiological requirements. To address this concept, we compared the IRE1-dependent gene expression programs of IRE1 knockout cells and IRE1-expressing cells driven by unfolded protein buildup (elicited by Tm) with those driven by ER translocation disruptions (elicited by RocA). We then corroborated the results in a different cell line, ISR^ON^ H4 cells, in the absence or presence of 4µ8C.

Expectedly, a targeted analysis of *XBP1* mRNA reads in our RNA sequencing datasets indicated the splicing of *XBP1* mRNA induced by both RocA and ISR^ON^ treatments, thus orthogonally validating the RT-PCR results described above (Fig. S7A – B).

Gene ontology (GO) term analyses of a subset of 1,495 mRNAs whose expression levels changed significantly (Abs log_2_FC > 0.5) upon RocA treatment in IRE1^WT^ cells compared to knockout controls revealed an enrichment of genes involved in translation and ER protein quality control. These processes included pathways that regulate protein synthesis, targeting and translocation, folding, glycosylation, and export (Fig. 4A, Fig. S8A). Similarly, comparing ISR^ON^ cells to 4µ8C-treated controls revealed an enrichment of the same processes (Fig. 4B, Fig. S8B). These analyses reveal a heretofore unrecognized IRE1-dependent gene expression program in response to co-translational translocation deficits.

**Figure 4.**
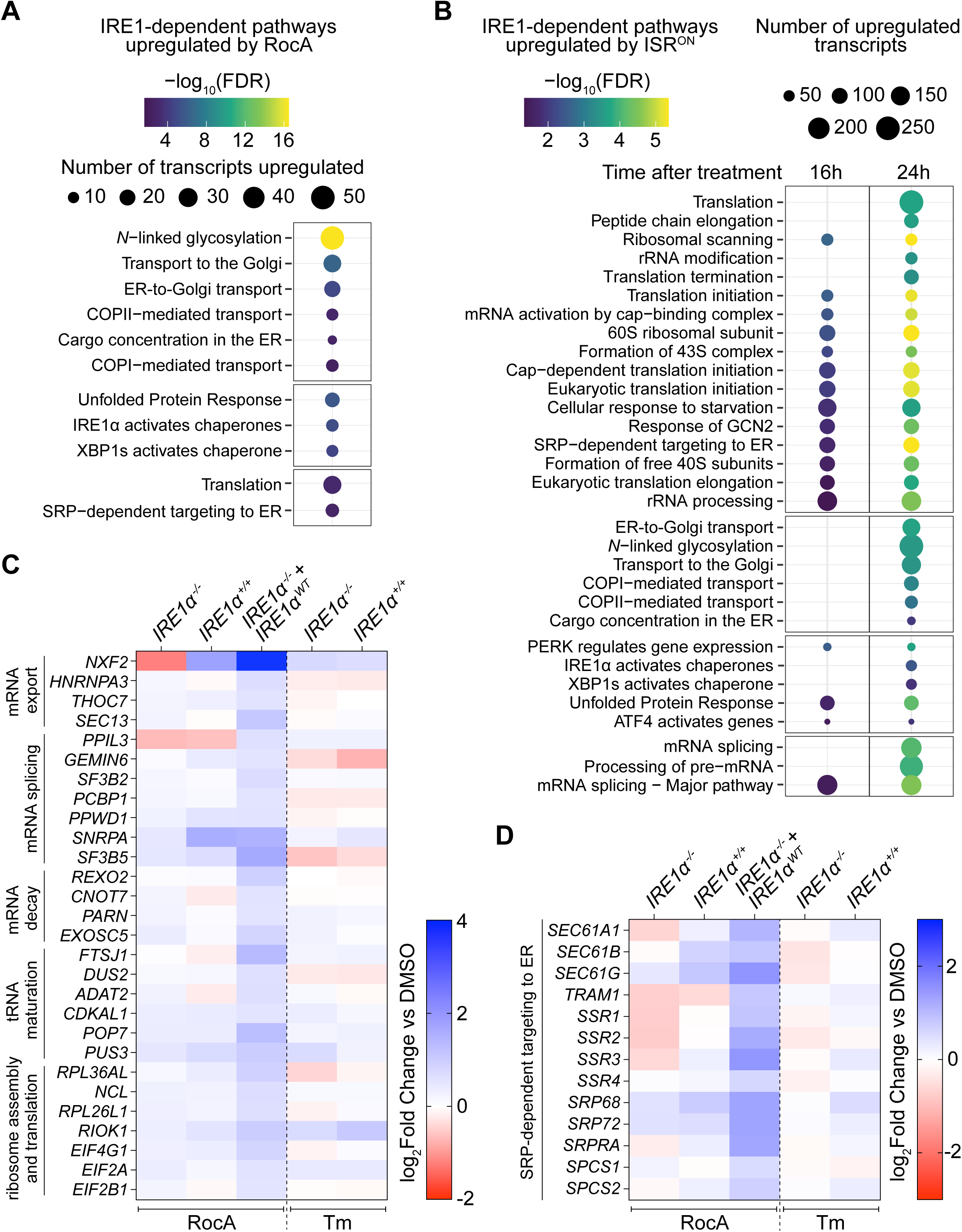
IRE1 controls distinct gene expression programs in response to reduced ER protein influx or increased unfolded protein buildup. (A-B) Bubble plots showing enriched Reactome pathways to which IRE1-dependent transcripts cluster as upregulated in (A) U-2 OS *IRE1α^-/-^* cells expressing mNeonGreen-tagged IRE1ɑ^WT^ upon addition of 5 ng/mL Dox and treated with 100 nM RocA for 16 hours (log_2_ FoldChange > 0.5 compared to DMSO controls), and (B) H4 cells with the activation of FKBP-PKR for 16 hours and 24 hours (continuous enrichment test as compared to controls). (C-D) Heatmaps showing log_2_FoldChanges of specific transcripts in U-2 OS *IRE1α^-/-^*cells, U-2 OS *IRE1α^+/+^* cells, and U-2 OS *IRE1α^-/-^* cells expressing mNeonGreen-tagged IRE1ɑ^WT^ upon addition of 5 ng/mL Dox. Cells were treated with 100 nM RocA for 16 hours as indicated. In parallel, U-2 OS *IRE1α^-/-^* cells and U-2 OS *IRE1α^+/+^* cells were treated with Tm for 4 hours. All log_2_FoldChanges were calculated in comparison to DMSO controls. The Reactome pathways to which the protein products of the differentially expressed transcripts belong are highlighted on the left of each panel.

To ascertain that this IRE1-dependent gene expression program is distinct from the IRE1 program induced by the UPR, we compared the expression changes of mRNAs induced by RocA and dependent on IRE1 to those induced by Tm, focusing on representative mRNAs in the prevalent GO categories. These analyses revealed unique TRES- or UPR-specific gene expression programs in addition to a common TRES/UPR gene expression program. The TRES-specific IRE1 program regulates genes involved in post-transcriptional gene expression, including those encoding components of mRNA splicing, maturation, export and decay, as well as ribosome assembly (Fig. 4C).

Notably, the TRES program also induced genes involved in regulating SRP-dependent co-translational targeting and translocation, including those encoding translocon subunits (*SEC61A1*, *SEC61B*, *SEC61G*), signal sequence receptor (*SSR1/2/3/4*), SRP (*SRP68/72*), SRP receptor (*SRPRA*), signal peptidase (*SPCS1/2*) and the translocation-associated membrane protein *TRAM1* (Fig. 4D). Conversely, the common TRES/UPR gene expression program includes genes encoding chaperones, foldases, trafficking machinery, and ERAD components (e.g., *HSPA5*, *HSP90B1*, *HYOU1*, *FICD*, *SYVN1*, *SEC22B*, *SEC24D*, *EDEM1*, *DERL2*) (Fig. S8C). Notably, the activation of this common program was exclusively dependent on IRE1 during TRES as several of these genes are known XBP1s targets. During the UPR, however, IRE1 signaling was largely dispensable for the upregulation of *RAB33B*, *HSP90B1*, *HYOU1*, *MANF* and *HSPA5* (Fig. S8C).

These data suggest that IRE1 controls specific homeostatic programs that prepare the ER and the secretory pathway to match particular physiological needs. These programs address fluctuations in ER function resulting from upsurges in ER protein-folding demands (UPR) or a reduction in ER protein influx upon co-translational translocation deficits (TRES).

## Discussion

IRE1 is known for its two detection and response modes to deviations from ER homeostasis that are triggered by ER protein folding perturbations and lipid bilayer stress, respectively. Here, we describe a hitherto third mode for IRE1 activation dubbed TRES: the sensing and responding to co-translational translocation deficits. Despite lowering the ER’s protein-folding burden akin to RIDD and RIDDLE, disrupted co-translational translocation and the subsequent reduction in ER protein influx imbalances the ER’s protein folding load and capacity. Fluctuations in protein synthesis, and consequently, oscillations in co-translational translocation occur, for example, during the maturation of B cells into antibody-producing plasma cells (*62*) and during normal functions of the endocrine pancreas, which are notably mediated by the ISR (*63*). Indeed, IRE1 and a primary product resulting from its activation, XBP1s, are essential to maintain the optimal secretory function of these cell types (*32*, *64*, *65*). We propose that a properly balanced ER function is defined by a “Goldilocks zone” in which homeostasis is optimized to the physiological needs of the cell. Displaying a U-shaped response to imbalances on either side, IRE1 emerges as the key ER homeostasis sentinel (Fig. 5, inset).

**Figure 5.**
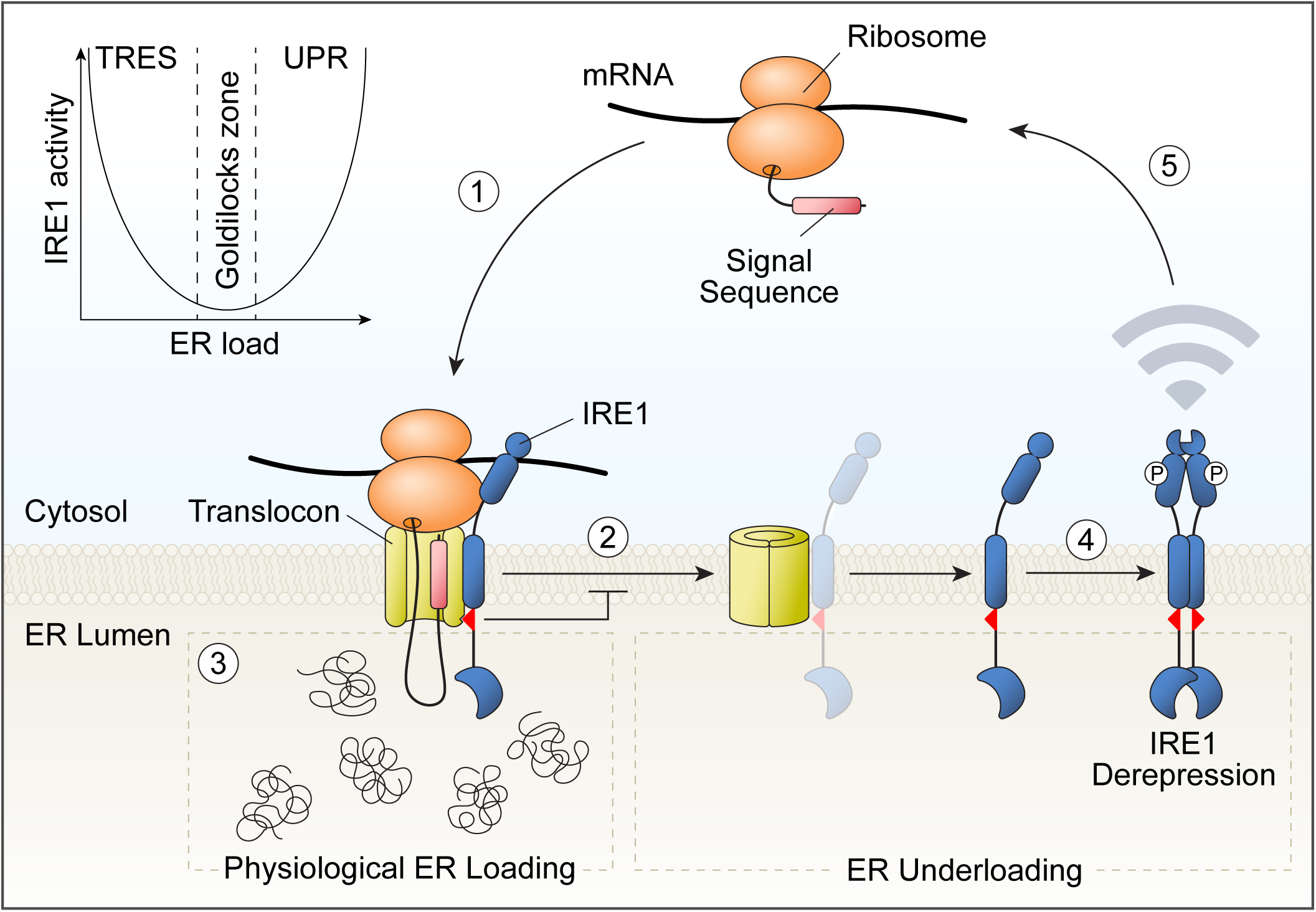
Model for IRE1 activation by reduced ER protein influx. *Schematic model*: (1) Co-translational translocation of signal-peptide bearing proteins into the ER facilitates IRE1’s association with the SEC61 translocon. These IRE1 species likely co-exist in a dynamic equilibrium. (2) The IRE1-SEC61 complex functions as an internal limiter of IRE1 activity when the ER protein load is optimal (the ‘Goldilocks zone’ (3) of ER protein load). (4) During TRES, disrupted co-translational translocation and reduced ER protein influx result in the closure of the SEC61 channel and the subsequent eviction of IRE1 from the translocon complex, leading to IRE1’s derepression and TRES signaling. (5) IRE1-driven gene expression programs in-turn prepare the cell for increased rates of protein synthesis and translocation into the ER by upregulating components of the co-translational translocation machinery. *Inset:* U-shaped response of IRE1 activity as a function of ER protein load.

### IRE1 follows distinct activation modes in response to ER stress resulting from ER overloading and underloading

Our observations support distinct IRE1 activation modes in response to fluctuations in ER load versus capacity, i.e., deviations from homeostasis. In one activation mode, increases in ER protein load cause BiP to dissociate from IRE1, enabling unfolded protein ligands in the ER lumen to bind to IRE1’s lumenal domain. Unfolded protein accumulation in the ER lumen promotes IRE1 oligomerization without dissociation from the translocon (see Fig. 1E, compares lanes 2 and 3), possibly by templated and spatially restricted IRE1 self-association. In contrast to this model, it has been proposed that IRE1’s association with the translocon regulates the attenuation of its signaling output (*10*, *48*). How IRE1 escapes attenuation imposed by SEC61 binding during unfolded protein stress remains an unresolved question.

During TRES, co-translational translocation deficiencies result in empty translocons that are conformationally distinct from those in an open state (*66–69*). We propose that the closure of the SEC61 channel evicts, and consequentially, derepresses IRE1 (Fig. 5). Our observations that disrupting co-translational translocation by inhibiting the channel, blocking translation initiation, or depleting SRP, induce *XBP1* mRNA splicing bolster this interpretation (Figs.1-3) and support a brake-release model for IRE1 activation upon dissociation from the translocon. Our findings are also consistent with earlier observations, indicating that depleting SEC61 translocon subunits activates IRE1 (*12*).

### IRE1 activation by TRES is unconventional and defined by specific characteristics

IRE1 activation in response to co-translational translocation defects has three unique characteristics: (i) it is independent of IRE1’s conventional ER stress sensing capabilities (Figs. 1-3), (ii) it sidesteps formation of large high-order oligomers visible as foci (Figs. S3, S4H, S6M), and (iii) it is muted in proportion to stress elicited by unfolded protein accumulation (Figs. 1C-D, 2D, 2F - compare lanes 3 and 5, 3K). TRES does not require IRE1’s lumenal domain or its transmembrane stress sensing domain. However, the lumenal domain amplifies signaling during TRES, perhaps due to the increased oligomerization potential of the intact protein. TRES activation of IRE1 does not result in formation of visible foci and is lower in magnitude compared to that observed in Tm-induced ER stress. Therefore, substantial *XBP1* mRNA splicing activity can result in the absence of visible foci. It is possible that as stress progresses to an irresolvable state (as would occur with Tm), continued IRE1 self-association leads to the larger assemblies that may be a preamble for IRE1 disposal by degradative means.

The proportionally muted response of TRES suggests IRE1 is less sensitive to co-translational translocation defects than to unfolded protein accumulation. The vast excess of translocons over IRE1 (back-of-the-envelope calculations indicated by ∼2,000-fold (*70*)) implies that IRE1 may sample a specialized subpopulation of translocons dedicated to challenging ER clients. This notion is supported by our previous finding that IRE1 preferentially associates with mRNAs encoding transmembrane proteins (*9*). Distinct translocon-associated factors could confer such specificity (*71*, *72*).

### IRE1 regulates different outputs in response to ER overload or underload

The distinct IRE1-dependent gene expression programs induced by UPR or TRES activation (Fig. 4) suggest that IRE1 activation modes are tuned to distinct physiological needs to reestablish homeostasis. The tuning can be achieved by combinatorial regulation via transcription factors. The UPR involves cooperation between XBP1s and ATF6-N (the cleaved nuclear form of ATF6, containing its transcription factor moiety) to induce gene expression programs (*27*, *60*) that mitigate ER overload. Conversely, TRES rebalances an underloaded ER by inducing XBP1s alone. ATF6/XBP1s heterodimers and XBP1s homodimers bind to distinct *cis*-regulatory elements, resulting in varied gene expression programs (*29*, *73*). In support of this notion, depleting SEC61 components, SRP machinery, eIF2 subunits, and eIF2B subunits—all of which disrupt co-translational translocation and hinder ER protein import—activates an XBP1s transcriptional reporter built with XBP1s-responsive *cis*-acting elements (*12*). It is also possible that XBP1s partners with other, yet-to-be-identified, transcription factors induced by TRES to induce a tailored gene expression program.

### TRES is integral to the ISR

TRES links IRE1 activity to the cell-wide attenuation of protein synthesis resulting from the diminished translation initiation that occurs upon eIF2 phosphorylation. Thus, TRES may fulfill an anticipatory role: IRE1 activation downstream of the ISR may prepare the cell for an incoming flux of secretory cargo once the ISR terminates and protein synthesis is restored. Other processes leading to decreased protein synthesis do not have the same impact on IRE1 activation, as indicated by experiments with the elongation inhibitors CHX and EME (Fig. S5C), which freeze ribosomes on their respective mRNAs and hence do not empty translocons. In this case, stalled ribosomes docked onto occupied translocons do not lead to IRE1 eviction from the translocon, preventing its activation. Thus, translational regulation by other means that stabilize existing polysomes would not engage TRES, while we predict that mechanisms that destabilize polysomes will activate it.

### TRES has pathophysiological implications

Mutations in *SEC63*, which encodes a protein that associates with the translocon, are linked to autosomal dominant polycystic liver disease (ADPLD), a genetic disease characterized by the formation of hepatic cysts, causing organ failure (*74*). Deletion of *Sec63* in mice, which recapitulates cyst formation in the kidney and liver, selectively activates IRE1 (*75*). In support of our findings on the homeostatic nature of IRE1 activation by TRES, *Xbp1* depletion in these mice exacerbates cyst formation while *Xbp1s* overexpression mitigates the disease (*75*). Moreover, IRE1 activation by TRES may have a role in oncological indications. For example, a SEC61 inhibitor (KZR-261) and Zotatifin are currently in clinical trials for advanced malignant solid tumors (NCT05047536 and NCT04092673), and Zotatifin is in clinical trials for estrogen receptor–positive HER2–negative breast cancer (NCT05101564) and low-grade serous ovarian cancer (NCT03675893). Considering IRE1’s adaptive roles and the dependence of some cancers on IRE1 activity (*7*), our results may inform on the therapeutic efficacy (or lack thereof) of targeting co-translational translocation in conjunction with IRE1 modulation to treat cancers. Importantly, given that dysregulation of the ISR is prominently linked to neurodegeneration (*38*), and the ISR inhibitors fosigotifator and DNL343 are in clinical trials (NCT05757141; NCT04948645; NCT06618118; NCT04297683; NCT05842941), it remains to be determined whether IRE1 activity is modulated in these conditions, and if targeting it would have additional therapeutic benefits. Taken together, the discovery of a third modality of IRE1 activation significantly expands our understanding of ER stress in physiology and pathology.

## Acknowledgments

We thank the members of the Acosta-Alvear, Walter, and Costa-Mattioli labs for insightful scientific discussions. We thank Dr. Ramanujan Hegde for advice on using SEC61-IN-1, Dr. Pascal Egea for the structural models of SEC61-IN-1 and CADA bound to the SEC61 translocon, and Dr. Rukmini Mukherjee for advice on the lipid bilayer stress experiments. We thank the Genomics Hub at Altos Labs BAI for their assistance with the RNA-sequencing experiments.

## Author contributions

Conceptualization: F.Z., A.S., P.W., D.A-A.

Methodology: F.Z., A.S., B.Y., T.L., J.W., R.Y., T.C., R.M., S.T., P.W., D.A-A. Investigation: F.Z., A.S., B.Y., J.E.C., T.C.

Writing - Original Draft: F.Z., A.S., P.W., D.A-A.

Writing - Review & Editing: All authors

Visualization: F.Z., A.S., B.Y., N.A.D., P.W., D.A-A.

Supervision: F.Z., A.S., S.T., M. C-M., P.W., D.A-A.

## Data, code, and materials availability

The data and materials generated in this study are available upon request through material and data transfer agreements with Altos Labs. The RNA sequencing datasets have been deposited to the Gene Expression Omnibus (GEO) server (accession number: GSE323967). All data are available in the main text or the supplementary materials.

## Supplementary Materials

Figs. S1 to S8

Materials and Methods

## Supplementary Materials

### Supplementary Figure Legends

**Figure S1.**
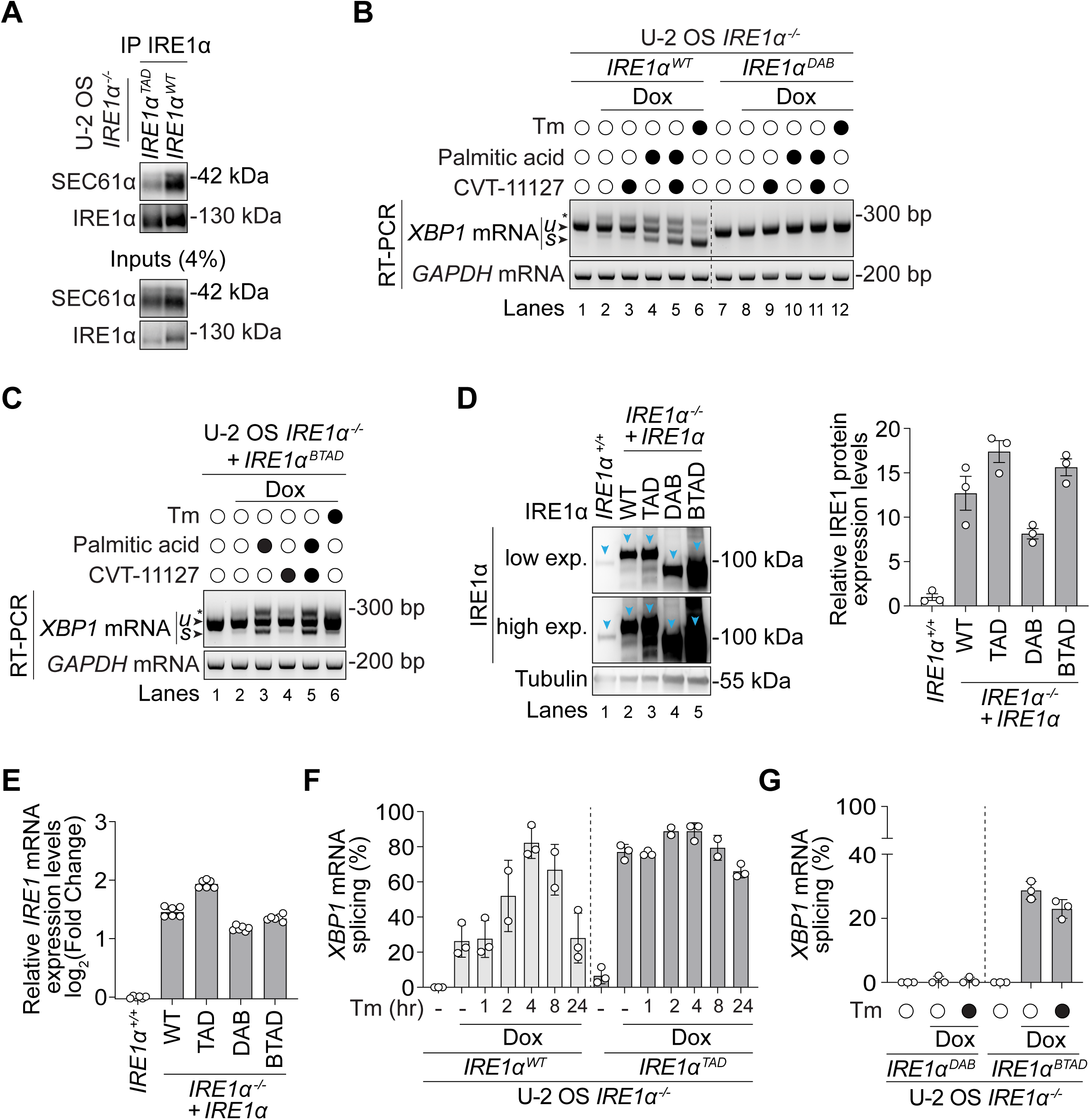
Functional characterization of IRE1 mutants used in this study. (A) Immunoprecipitation of mNeonGreen-tagged IRE1α^WT^ and IRE1α^TAD^ in lysates obtained from U-2 OS *IRE1α^-/-^*cells expressing the respective constructs after the addition of 5 ng/mL Dox for 48 hours. Immunoblotting was performed using antibodies against SEC61α and IRE1α. (B-C) RT-PCR of *XBP1* and *GAPDH* mRNAs in lysates obtained from U-2 OS *IRE1α^-/-^* cells expressing mNeonGreen-tagged IRE1α^WT^ and IRE1α^DAB^ (B) or mNeonGreen-tagged IRE1^BTAD^ (C) upon addition of 5 ng/mL doxycycline (Dox). Cells were treated with 5 µg/mL Tm for 4 hours, 500 µM palmitic acid for 6 hours, or 1 µM CVT-11127 for 24 hours as indicated. u - unspliced; s - spliced *XBP1* mRNA amplicons, respectively. The asterisk indicates a hybrid amplicon generated by annealing of one strand of *XBP1*u and one strand of *XBP1*s during the PCR. GAPDH: loading control. Data are representative of two independent experiments. (D) Immunoblots of IRE1α and tubulin in lysates obtained from U-2 OS (*IRE1α^+/+^*) cells and U-2 OS *IRE1α^-/-^* cells expressing mNeonGreen-tagged IRE1α^WT^, IRE1α^TAD^, IRE1α^DAB^, and IRE1α^BTAD^ upon addition of 5 ng/mL doxycycline (Dox) for 48 hours (left). Relative protein expression levels of IRE1 were normalized to tubulin and quantified as a fold change of the mean ± SEM over endogenous IRE1α levels (right) (N=3). (E) RT-qPCR of *IRE1α* mRNA levels in lysates obtained from U-2 OS (*IRE1α^+/+^*) cells and U-2 OS *IRE1α^-/-^*cells expressing mNeonGreen-tagged IRE1α^WT^, IRE1α^TAD^, IRE1α^DAB^, and IRE1α^BTAD^ upon addition of 5 ng/mL doxycycline (Dox) for 48 hours. Relative *IRE1α* mRNA expression levels were normalized internally to the reference gene *GAPDH* and expressed as log_2_(fold change) over the levels in U-2 OS (*IRE1α^+/+^*) cells. Data: mean ± SEM (N=6). (F-G) Quantification of *XBP1* mRNA splicing from experiments shown in Fig. 1B (F) and Fig. 1C (G). Data: mean ± SD (N ≥ 2).

**Figure S2.**
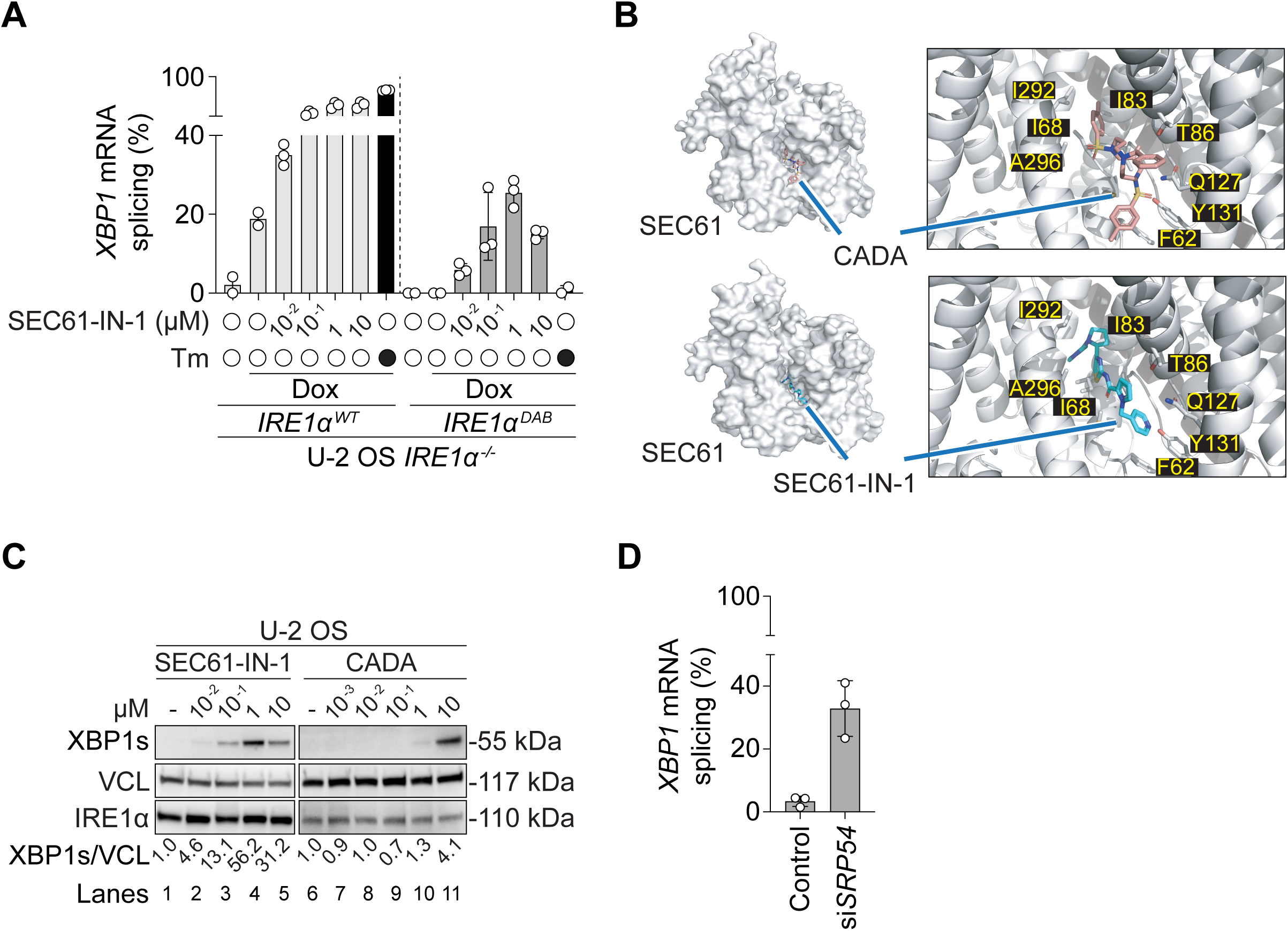
Inhibition of co-translational translocation activates IRE1. (A) Quantification of *XBP1* mRNA splicing from experiments shown in Fig. 1D. Data: mean ± SD (N ≥ 2). (B) Molecular docking of SEC61-IN-1 into the structure of the SEC61 translocon, and Cryo-EM structure of CADA bound to the SEC61 translocon (PDB: 8DO2). Panels on the right show the zoomed in views of the small molecules occupying the central pore of SEC61α. (C) Immunoblots of XBP1s, Vinculin (VCL) and IRE1α in lysates obtained from U-2 OS cells treated with CADA and SEC61-IN-1 for 16 hours at the indicated concentrations. The quantification of the proportion of spliced XBP1 protein is indicated below. Immunoblots for each treatment were processed in parallel on different gels. Data are representative of two independent experiments. (D) Quantification of *XBP1* mRNA splicing from experiments shown in Fig. 1G. Data: mean ± SD (N ≥ 2).

**Figure S3.**
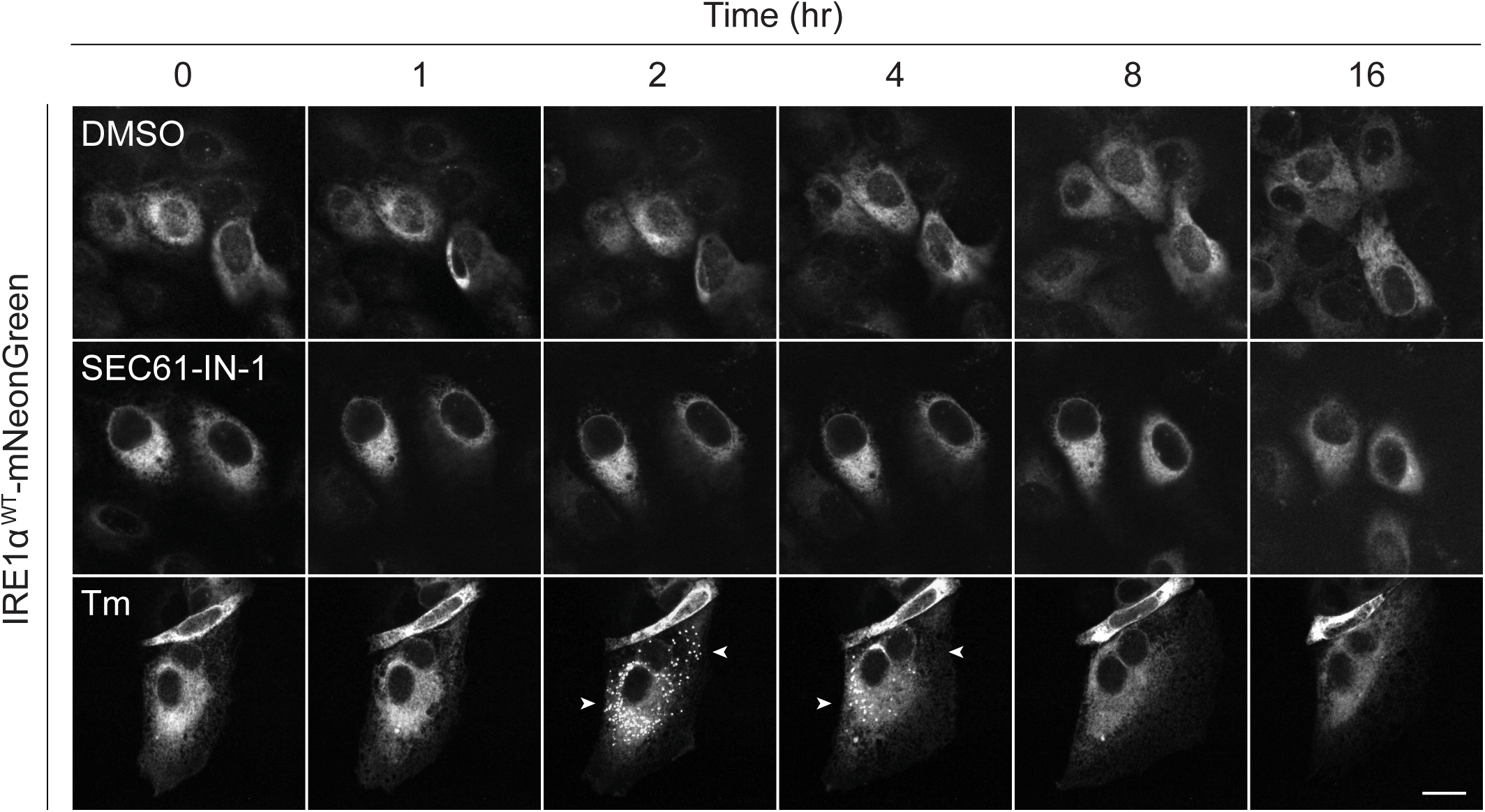
SEC61 translocon inhibition does not induce large IRE1 foci. Representative micrographs of U-2 OS *IRE1α*^-/-^ cells expressing mNeonGreen-tagged IRE1α^WT^ upon addition of 5 ng/mL doxycycline and treated with 1 µM SEC61-IN-1 or 5 µg/mL Tm for the indicated times. Arrowheads indicate high-order mNeonGreen-tagged IRE1α foci observable by confocal fluorescence microscopy. Scale bar = 10 µm.

**Figure S4.**
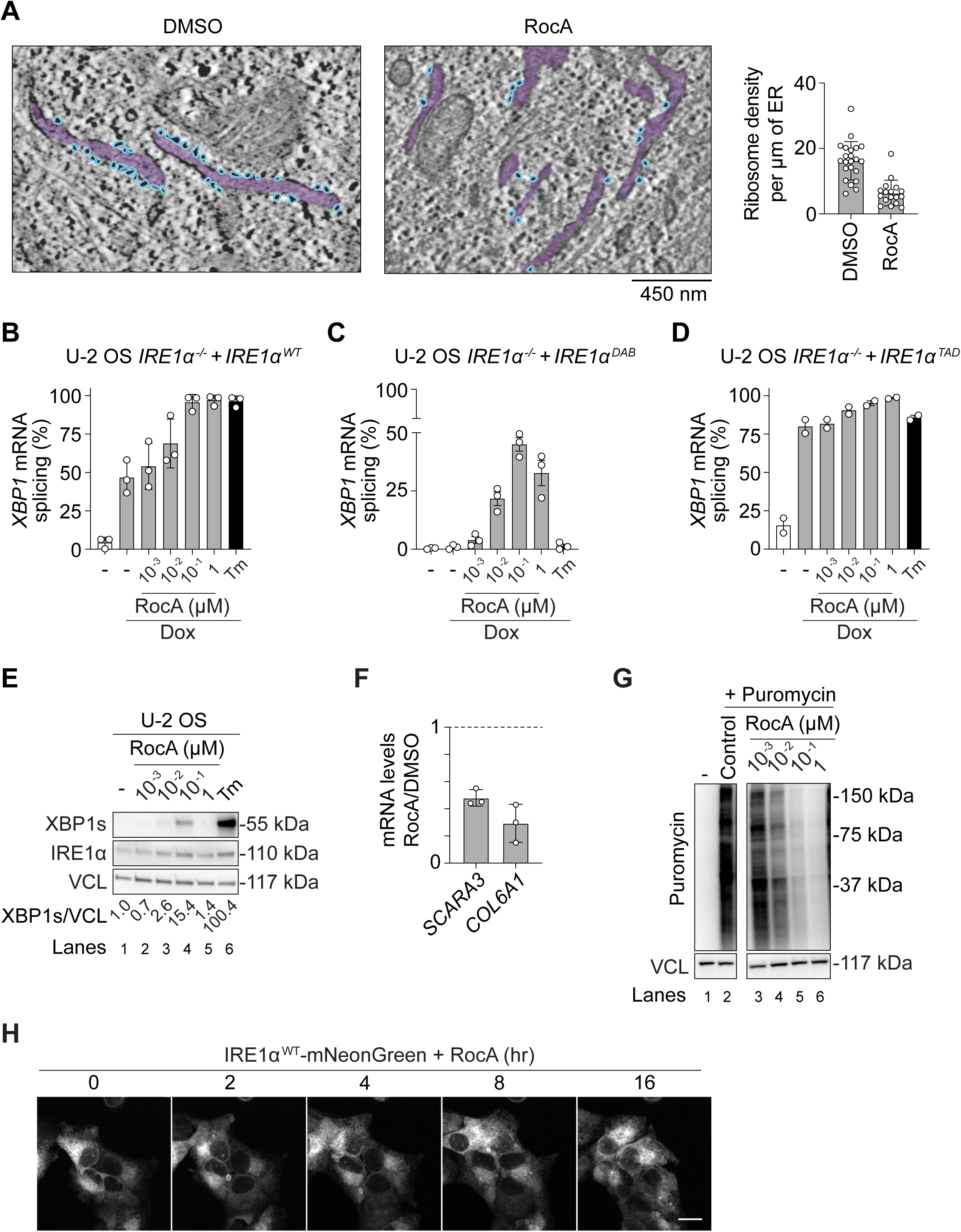
Inhibiting translation initiation activates IRE1. (A) Quantification of ribosome density based on analysis of transmission electron micrographs of ER membranes (shaded in purple) in U-2 OS cells treated with 100 nM RocA for 16 hours. Electron dense ribosome particles are highlighted in blue. Data: Mean ± SEM (N=22 micrographs for control and N=17 micrographs for RocA). (B-D) Quantification of *XBP1* mRNA splicing from experiments shown in Fig. 2B, 2D and 2E. Data: mean ± SD (N ≥ 2). (E) Immunoblots of XBP1s, IRE1α, and Vinculin (VCL) in lysates obtained from U-2 OS cells treated with RocA for 16 hours at the indicated concentrations or 5 µg/mL Tm for 4 hours. The quantification of the proportion of spliced XBP1 protein is indicated below the immunoblots. Data are representative of three independent experiments. (F) RT-qPCR of *SCARA3* and *COL6A1* mRNAs in H4 cells treated with 1 µM RocA for 16 hours. Expression levels were normalized internally to 28S ribosomal RNA. Values are expressed as fold change over the levels in H4 cells treated with DMSO (indicated with a dashed line). Data: Mean ± SEM (N = 3). (G) U-2 OS cells treated with RocA at the indicated concentrations for 8 hours were subjected to puromycin pulse-chase analyses to assess nascent protein synthesis rates. Immunoblotting of lysates was performed using antibodies to puromycin and VCL. Data are representative of two independent experiments. Lanes were cropped from the same gel. (H) Confocal micrographs of U-2 OS *IRE1α^-/-^*cells expressing mNeonGreen-tagged IRE1α^WT^ upon addition of 5 ng/mL Dox and treated with 100 nM RocA for the indicated times. Data are representative of two independent experiments. Scale bar = 10 µm.

**Figure S5.**
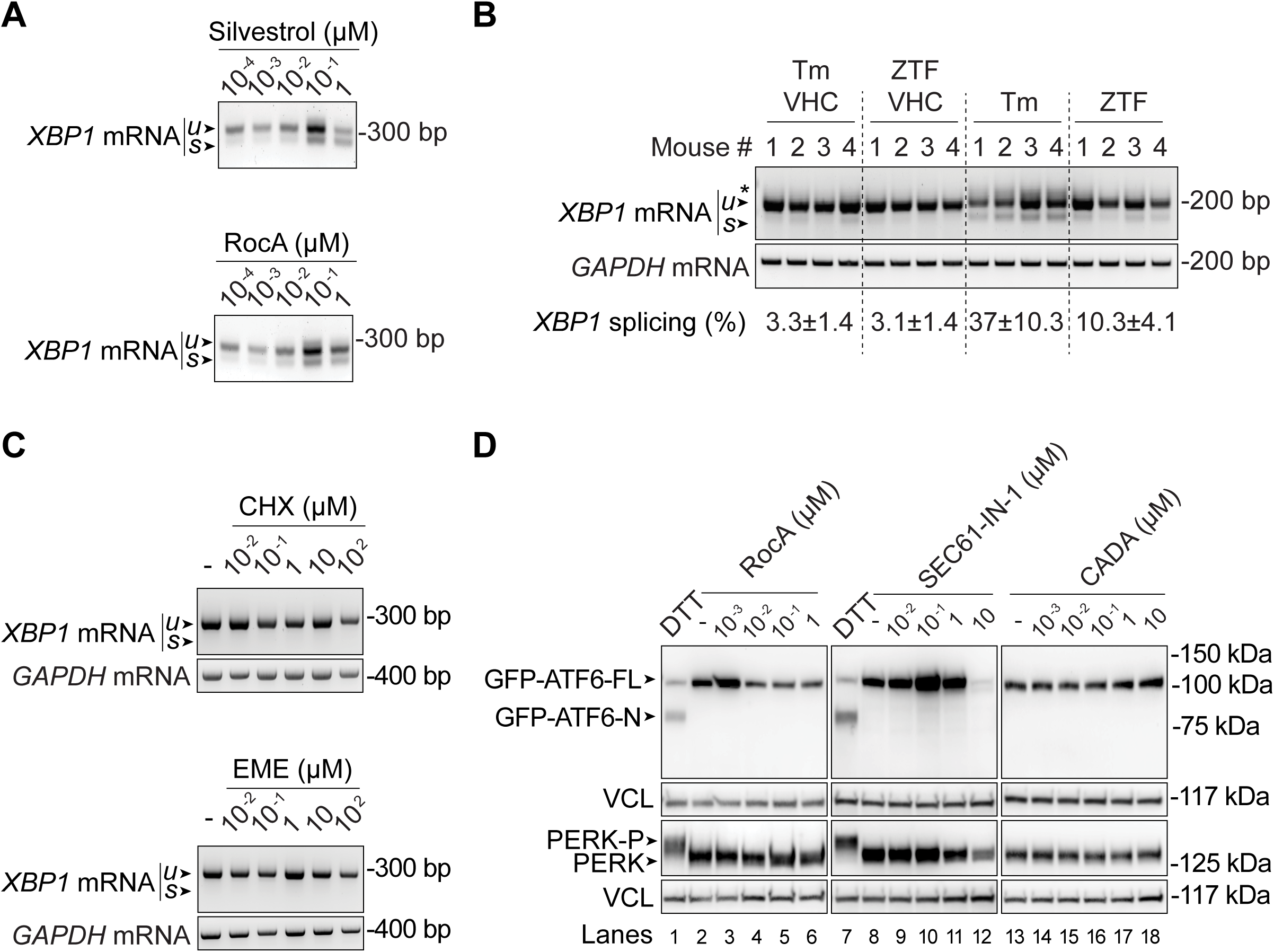
Inhibiting co-translational translocation activates IRE1 but not ATF6 or PERK. (A) RT-PCR of *XBP1* mRNA in lysates obtained from H4 cells treated with RocA and silvestrol at the indicated concentrations for 16 hours. u/s unspliced and spliced *XBP1* amplicons, respectively. (B) RT-PCR of *XBP1* and *GAPDH* mRNAs in lysates obtained from liver tissues of mice treated with 1 mg/kg ZTF or 1 mg/kg Tm for 16 hours. *GAPDH*: loading control. The percentage of *XBP1* splicing is shown below the RT-PCR gels and represents Mean ± SEM of N = 8 mice per treatment. VHC - vehicle control. (C) RT-PCR of *XBP1* and *GAPDH* mRNAs in lysates obtained from U-2 OS cells treated with cycloheximide (CHX) and emetine (EME) at the indicated concentrations for 16 hours. *GAPDH*: loading control. (D) Immunoblots of GFP-ATF6 in lysates obtained from U-2 OS cells stably expressing ATF6-GFP (top) and PERK in lysates obtained from U-2 OS cells (bottom). Cells were treated with RocA, SEC61-IN-1 and CADA at the indicated concentrations for 16 hours, or 1 mM DTT for 1 hour. GFP-ATF6-N: cleaved ATF6 N-terminus. PERK-P: phosphorylated PERK. VCL: loading control. Immunoblots for each treatment were processed in parallel on different gels. Data are representative of two independent experiments.

**Figure S6.**
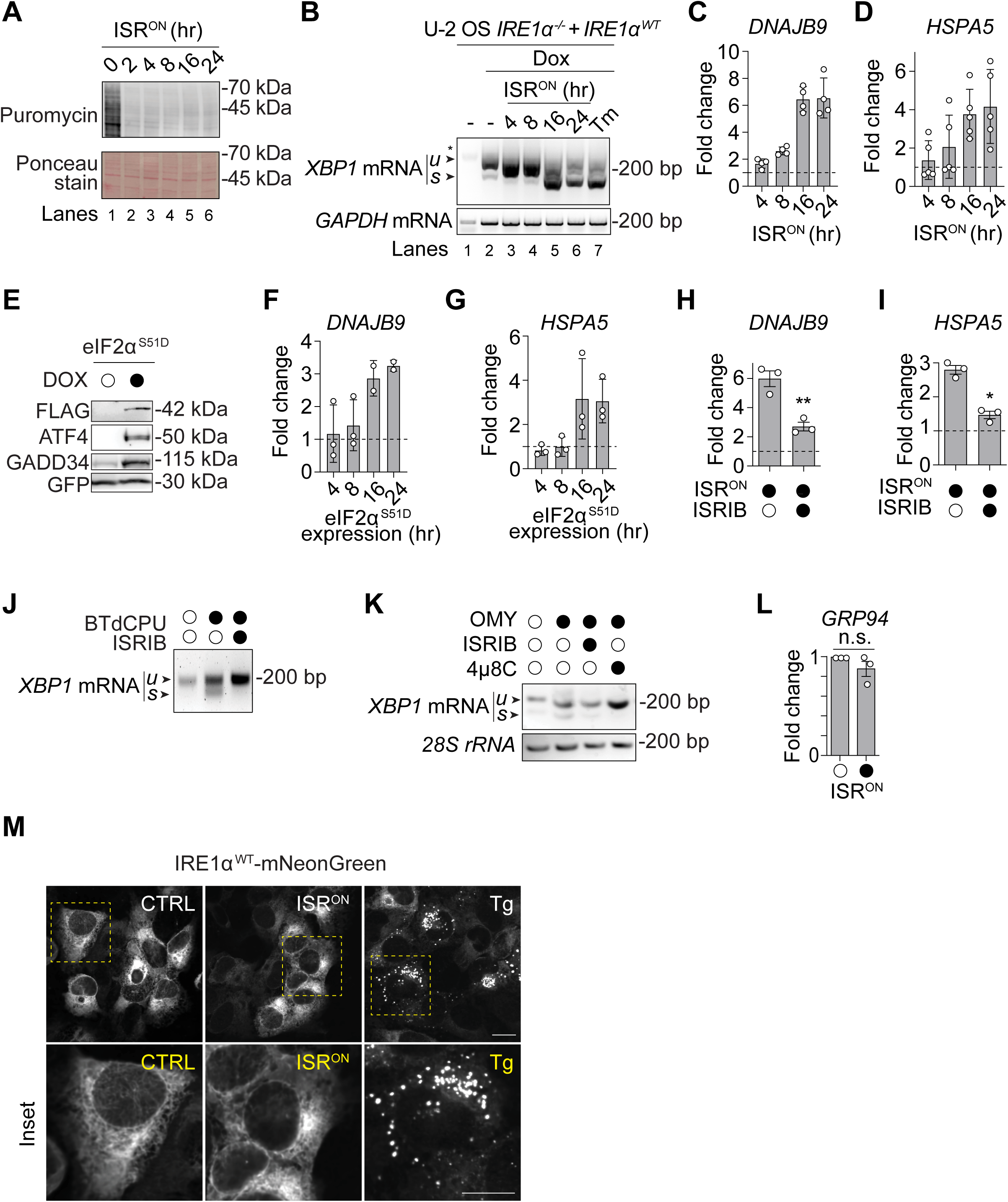
ISR induction activates IRE1. (A) H4 cells expressing FKBP-PKR and treated with the homodimerizer for the indicated times were subjected to puromycin pulse-chase analyses to assess nascent protein synthesis rates. Immunoblotting was performed using antibodies to puromycin. Ponceau staining: loading control. Data are representative of three independent experiments. (B) RT-PCR of *XBP1* and *GAPDH* mRNAs in U-2 OS *IRE1α^-/-^* cells expressing FKBP-PKR and mNeonGreen-tagged IRE1α^WT^. IRE1α^WT^ expression was induced with 20 ng/mL doxycycline (Dox). Cells were treated with 5 µg/mL Tm for 4 hours or the homodimerizer for the indicated times. Data are representative of three independent experiments. (C-D) RT-qPCR of *DNAJB9* (C) and *HSPA5* (D) mRNAs in H4 cells expressing FKBP-PKR and treated with the homodimerizer. Data: Mean ± SEM of fold changes normalized to the levels in untreated controls; N = 4 for (C), and N = 5 for (D). (E) Immunoblots of FLAG-tagged eIF2α^S51D^, ATF4, GADD34, and GFP in lysates obtained from H4 cells expressing eIF2α^S51D^ hosted in a construct that also expresses GFP for 24 hours. GFP: loading control. (F-G) RT-qPCR of *DNAJB9* (F) and *HSPA5* (G) mRNAs in H4 cells expressing eIF2α^S51D^ hosted in a construct that also expresses GFP for the indicated times. Data: Mean ± SEM of fold changes normalized to the levels in untreated controls (N ≥ 2). (H-I) RT-qPCR of *DNAJB9* (H) and *HSPA5* (I) mRNAs in H4 cells expressing FKBP-PKR and treated with the dimerizer, with or without co-treatment with 800 nM ISRIB, for 16 hours. Data: Mean ± SEM of fold changes normalized to the levels in untreated controls (N = 3). *p-value < 0.05, **p-value < 0.01, Student’s t-test. (J) RT-PCR of *XBP1* mRNA in H4 cells treated with 10 µM BtdCPU with or without co-treatment with 800 nM ISRIB for 24 hours. Data are representative of three independent experiments. (K) RT-PCR of *XBP1* mRNA and *28S* ribosomal RNA in HEK-293 cells treated with 5 ng/µl (6.3 µM) oligomycin (OMY) with or without co-treatment with either 10 µM 4µ8C or 800 nM ISRIB for 16 hours. Data are representative of three independent experiments. (L) RT-qPCR of *GRP94* mRNA in H4 cells expressing FKBP-PKR and treated with the homodimerizer for 16 hours. Data: Mean ± SEM of fold changes normalized to the levels in control cells. (M) Confocal micrographs of U-2 OS *IRE1α^-/-^*cells expressing FKBP-PKR and mNeonGreen-tagged IRE1α^WT^ and treated with 300 nM thapsigargin (Tg) for 4 hours or the homodimerizer for 16 hours (N = 3). Scale bar = 10µm.

**Figure S7.**
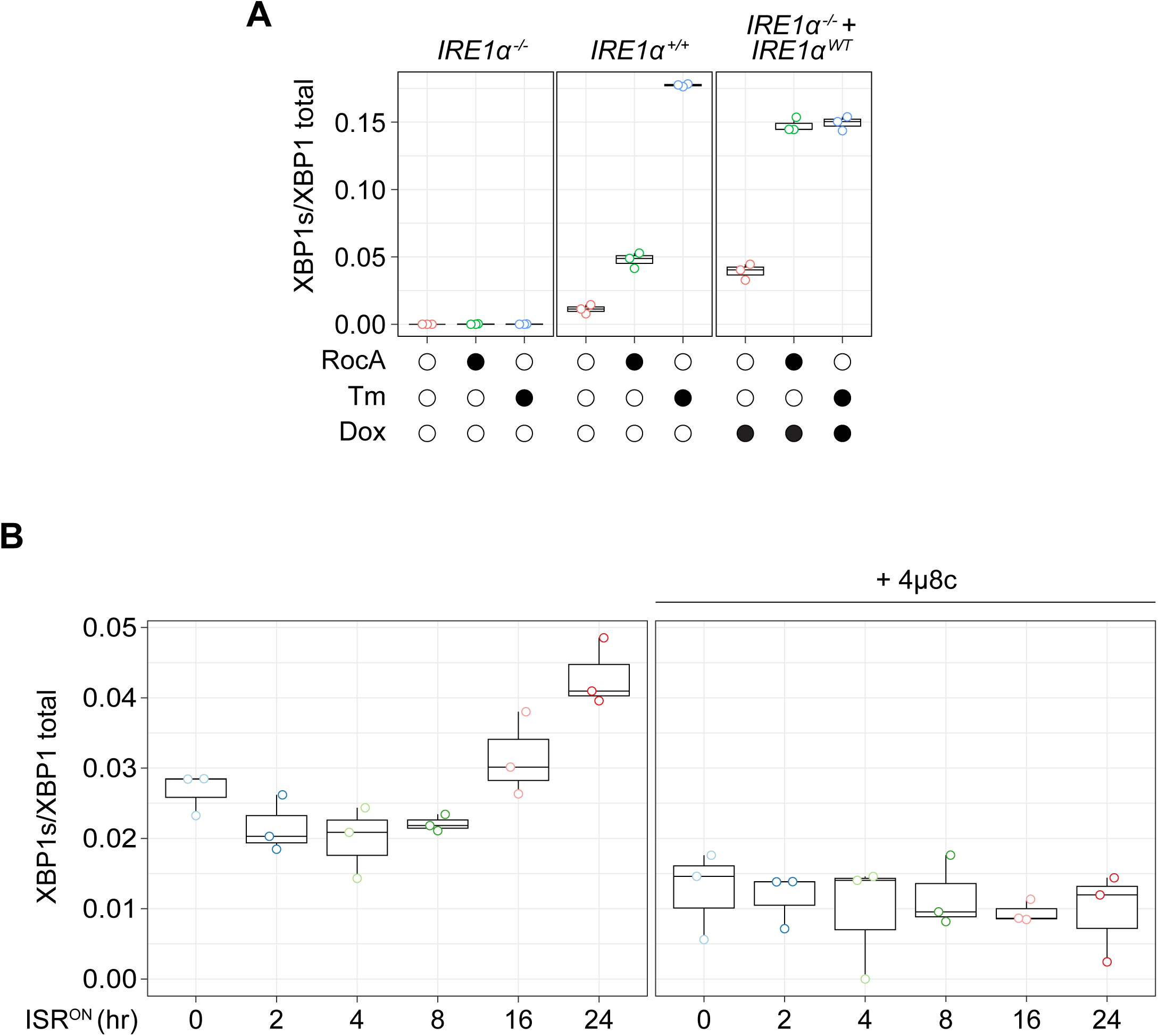
RNA sequencing reads validate *XBP1* splicing induction by RocA and ISR^ON^ treatments. (A-B) RNA-sequencing based validation of *XBP1* mRNA splicing in (A) U-2 OS *IRE1α*^-/-^ cells, U-2 OS *IRE1α^+/+^* cells, and U-2 OS *IRE1α*^-/-^ cells expressing mNeonGreen-tagged IRE1α^WT^ upon addition of 5 ng/mL Dox and treated with 100 nM RocA for 16 hours or Tm for 4 hours as indicated; and (B) H4 cells expressing FKBP-PKR and treated with the homodimerizer for the indicated times. Data: Proportion of *XBP1s* (Ratio of counts per million *XBP1* reads).

**Figure S8.**
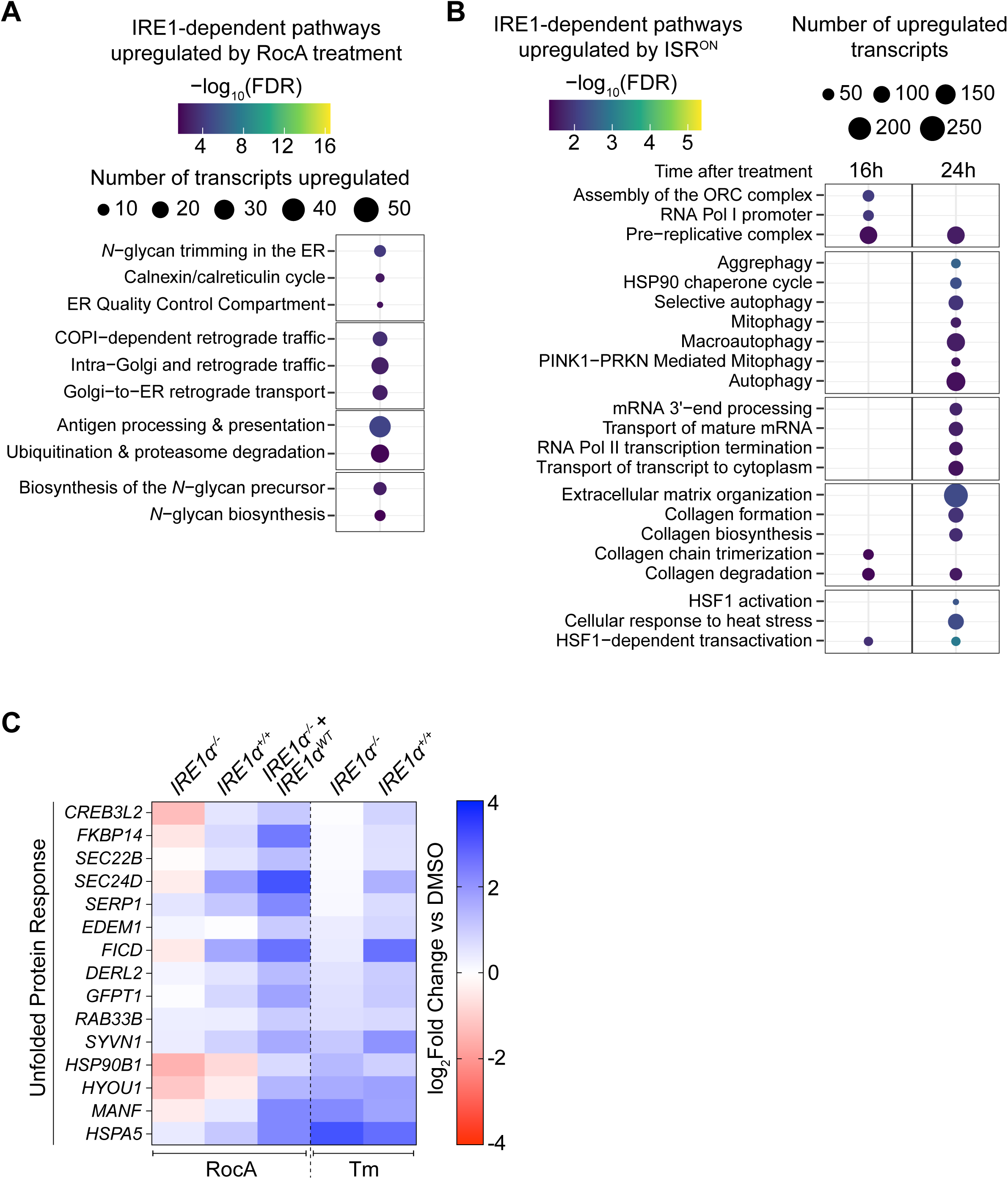
IRE1 regulates a common set of transcripts in response to RocA or tunicamycin treatments. (A-B) Bubble plots showing differentially enriched Reactome pathways to which IRE1-dependent transcripts cluster as upregulated in (A) U-2 OS *IRE1α*^-/-^ cells expressing mNeonGreen-tagged IRE1α^WT^ upon addition of 5 ng/mL Dox and treated with 100 nM RocA for 16 hours (log_2_ FoldChange > 0.5 compared to DMSO controls), and (B) H4 cells with the activation of FKBP-PKR for 16 hours and 24 hours (continuous enrichment test as compared to controls). (C) Heatmap showing log_2_ FoldChanges of common UPR transcripts in U-2 OS *IRE1α*^-/-^ cells, U-2 OS *IRE1α^+/+^*cells, and U-2 OS *IRE1α*^-/-^ cells expressing mNeonGreen-tagged IRE1α^WT^ upon addition of 5 ng/mL Dox. Cells were treated with 100 nM RocA for 16 hours. In parallel, U-2 OS *IRE1α*^-/-^ and U-2 OS *IRE1α^+/+^* cells were treated with Tm for 4 hours. All log_2_ FoldChanges are relative to DMSO controls.

## Materials and Methods

### Cell culture, genetic knockdown, transduction, and drug treatments

U-2 OS (ATCC HTB-96), H4 (ATCC HTB-148), and HEK-293 (ATCC CRL-1573) cells were cultured in Dulbecco’s Modified Eagle Medium (DMEM) supplemented with 10% fetal bovine serum (FBS) and 1X GlutaMax (Gibco). All cell lines used in the study tested negative for Mycoplasma contamination when assayed with the Universal Mycoplasma Detection Kit. The U-2 OS Flp-In *IRE1α*^-/-^ cells were generated as previously described (*41*). The Flp-In system was used to reconstitute U-2 OS *IRE1α^-/-^*cells with mNeonGreen-tagged IRE1α variants. The parental cells were plated in 6-well plates at a density of 17,000 cells/cm^2^. The following day, they were co-transfected with 1.7 μg of the Flp recombinase expression vector, pOG44 (Thermo Fisher V600520), and 300 ng of pcDNA5/FRT/TO encoding the tagged IRE1α variants (ThermoFisher V652020). Cells with successful integration were selected with 50 μg/mL hygromycin B (Thermo Fisher 10687010). IRE1α expression was induced by addition of doxycycline for a total duration of 36 - 48 hours unless otherwise stated. U-2 OS mNeonGreen-tagged *IRE1α*^-/-^ cells expressing IRE1α^WT^ or IRE1α^ΔI19-D408+W457A^ and FKBP-PKR-P2A-iRFP were obtained through lentiviral infection and were sorted using fluorescent-activated cell sorting (FACS). H4 FKBP-PKR and Tet-On eIF2α S51D-expressing cells were generated as previously described (*54*, *55*). H4 cells expressing the CRISPRi machinery (kind gift of Martin Kampman, UCSF) were used to establish the ATF6 KD cell line by transduction with VSV-G pseudotyped lentiviruses encoding three small guide RNAs. The sgRNA sequences were obtained from the human genome-scale CRISPRi library developed by the laboratory of Jonathan Weissman (MIT, Whitehead Institute). Seventy-two hours after infection, the cells were selected with 1 µg/ml of puromycin (Sigma-Alrich). After selection, a sub-clonal population of cells expressing the sgRNAs was sorted using FACS. Transient gene depletion was achieved through RNA interference (RNAi) by transfecting pooled siRNAs (Dharmacon) using Lipofectamine RNAi MAX (Invitrogen) according to the manufacturer’s recommendations. Subsequent analyses were conducted 72-96 hours after transfection. All drugs used in this study are listed in Table 1. Drug concentrations and treatment durations are indicated in the main text and figure legends.

**Table 1.** Drugs used in this study.

| Drug | Vendor | Catalogue number | Resuspension solvent | Concentration used (unless otherwise stated) |
| --- | --- | --- | --- | --- |
| SEC61-IN-1 (A317) | MCE | HY-139615 | DMSO (dimethyl sulfoxide) | 0.01-10 µM |
| Rocaglamide | MCE | HY-19356 | DMSO | 1-1000 nM |
| Cycloheximide | Arcos (Fisher Scientific) | AC357420010 | H<sub>2</sub>O | 0.01-100 µM |
| Silvestrol | Abcam | ab285419 (lot# GR3431087-2) | DMSO | 0.1-1000 nM |
| Zotatifin | MCE | HY-112163 | 5% Dextrose in H<sub>2</sub>O | 1 mg/kg |
| Puromycin | Thermo Fisher | AAJ61278MC | DMSO | 1 - 2 µg/mL |
| Cyclotriazadisulfonamide (CADA) | Sigma-Aldrich | 5343370001 | DMSO | 0.001-10 µM |
| AP-20187 (FKBP homodimerizer) | Takara | 635058 | Ethanol | 100 nM |
| Tunicamycin | Sigma-Aldrich | SML1287 | DMSO | 5 µg/mL<br>1 mg/kg |
| Thapsigargin | Adipogen Life Sciences | AG-CN2-0003 | DMSO | 300 nM |
| 4µ8C | SelleckChem | S7272 | DMSO | 30 µM |
| DTT (Dithiothreitol) | Sigma-Aldrich | D0632 | H<sub>2</sub>O | 1 mM |
| *Trans*-ISRIB | Torcis (Fisher Scientific) | 5284 | DMSO | 800 nM |
| Ceapin-A7 | Torcis (Fisher Scientific) | 6955/10 (69-551-0) | DMSO | 5 µM |
| Oligomycin | Sigma-Aldrich | O4876 | DMSO | 25 µM |
| BTdCPU | MCE | HY-118266 | DMSO | 10 µM |
| KIRA 8 | CME | 1630086-20-2 | DMSO | 1 µM |
| Doxycycline | Sigma-Aldrich | D9891 | H<sub>2</sub>O | 5 - 20 ng/mL |
| CVT-11127 | MCE | HY-113638 | DMSO | 1 μM |
| Sodium palmitate | Sigma-Aldrich | SLCN0277 | 0.1 M NaOH | 500 μM |

### Puromycylation assay

For the experiments in Fig. S4G, U-2 OS cells were treated with 2 µg/ml puromycin for 10 minutes, followed by a wash with 1X PBS. The cells were chased with complete medium with or without RocA for 50 minutes, washed with 1X PBS, lysed with RIPA buffer (1% Triton X-100, 20 mM Tris–HCl pH 8.0, 0.1% SDS, 0.05% sodium deoxycholate, 150 mM NaCl, 10 mM Na3VO4, 40 mM β-glycerophosphate and 10 mM NaF), and the lysates subjected to immunoblotting. For the experiments in Fig. S6A, H4 cells were grown to ∼70% confluency and incubated with 1 µg/ml puromycin for 20 minutes at 37 °C in complete media. After three washes with 1X PBS, the cells were lysed, and the samples were prepared for immunoblotting of puromycylated peptides.

### Transmission Electron Microscopy (TEM)

U-2 OS cells were grown on 35 mm plain-bottom MatTek dishes until they reached ∼60-70% confluency, at which point they were fixed in a solution of 3% glutaraldehyde and 3% PFA in 1xPBS at pH 7.4 (EMS product 16538-06). Samples were processed for TEM by the Electron Microscope Lab at UC Berkeley, following their standard protocol (https://em-lab.berkeley.edu/EML/tem-generic-protocol). Ultrathin (60 nm) and thick (300 nm) sections were cut by microtome and collected onto formvar-coated copper slot grids. Imaging was performed on a Krios G4 TEM (Thermo Fisher Scientific) operated at 300 kV, equipped with a Selectris X energy filter and a Falcon 4i detector (Thermo Fisher Scientific). Images were acquired at a pixel size of 1-2 nm. Tilt series were collected from -60 to +60 at 2 increments using Tomography 5 software (Thermo Fisher Scientific). The tilt series were processed and reconstructed using TomoLive (Thermo Fisher Scientific), and the 3D reconstructions were analyzed in Amira software (Thermo Fisher Scientific). ER lengths were manually traced and quantified in 2D tomographic slices, and ER-associated ribosomes were manually identified.

### *In vivo* studies

All animal procedures were approved by the Altos Labs Institutional Animal Care and Use Committee (IACUC) under protocol 21-105-001. All experiments were conducted strictly with the Guide for the Care and Use of Laboratory Animals (National Research Council, 8th edition) and relevant national regulations. Male C57BL/6J mice (The Jackson Laboratory, Bar Harbor, ME, USA; Strain #000664), 9 weeks of age, were used for the study. Animals were housed in individually ventilated cages with *ad libitum* access to a standard chow diet (Teklad Irradiated Global Soy Protein-Free Extruded Rodent Diet, Inotiv cat#2029X) and purified water. The housing environment was maintained on a 12-hour light/dark cycle (lights on at 07:00 AM, lights off at 07:00 PM), with controlled temperature (20-26 °C) and humidity (40-60%). Mice were acclimated to the facility for 1 week before the start of experimental procedures. Animals were randomly assigned to experimental groups. Experimenters were blinded to treatment groups during drug administration, sample collection, and subsequent data analysis. Tunicamycin (Sigma Aldrich Cat no. SML1287) was diluted 1:25 in 150 mM dextrose to achieve a final concentration of 200 µg/mL. Zotatifin (MCE HY-112163) was resuspended in 5% Dextrose in ddH_2_O at a final concentration of 200 µg/mL. The solutions were filter-sterilized using a 0.22 µm syringe filter. Animals received a single intraperitoneal (IP) injection of equal volumes of vehicle control or drugs as follows: Tunicamycin vehicle, 150 mM Dextrose in 4% DMSO; Zotatifin vehicle, 5% Dextrose (278 mM Dextrose) via IP injection. At 16 hours post-injection, mice were terminally anesthetized with isoflurane (2% for induction and 1.5% for maintenance), and their livers were collected after a 1X PBS perfusion. Livers were rinsed in ice-cold PBS and flash-frozen in liquid nitrogen before storage at -80 °C until further analysis.

### Immunoblotting

For experiments in Fig. 1D, 3C, 3E, 3I and S6A, cell lysates were collected directly in Laemmli SDS-PAGE sample buffer (62.5 mM Tris-HCl pH 6.8, 2% SDS, 10% glycerol, and 0.01% bromophenol blue). Lysates were briefly sonicated and supplemented with fresh 5% 2-mercaptoethanol before heat denaturation and separation by SDS-PAGE. Lysates were separated on SDS-PAGE gels and transferred onto nitrocellulose membranes for immunoblotting. Immunoreactive bands were detected by enhanced chemiluminescence using horseradish peroxidase (HRP)-conjugated secondary antibodies.

For all other experiments, the cells were washed three times with 1X PBS and immediately collected at 4 °C in RIPA lysis buffer containing phosphatase inhibitors (1% Triton X-100, 20 mM Tris–HCl pH 8.0, 0.1% SDS, 0.05% sodium deoxycholate, 150 mM NaCl, 10 mM Na_3_VO_4_, 40 mM β-glycerophosphate and 10 mM NaF) and complete protease inhibitors (Roche). The detergent-soluble supernatant fractions were immediately processed for SDS-PAGE and immunoblotting. Antibodies are listed in Table 2.

**Table 2.**
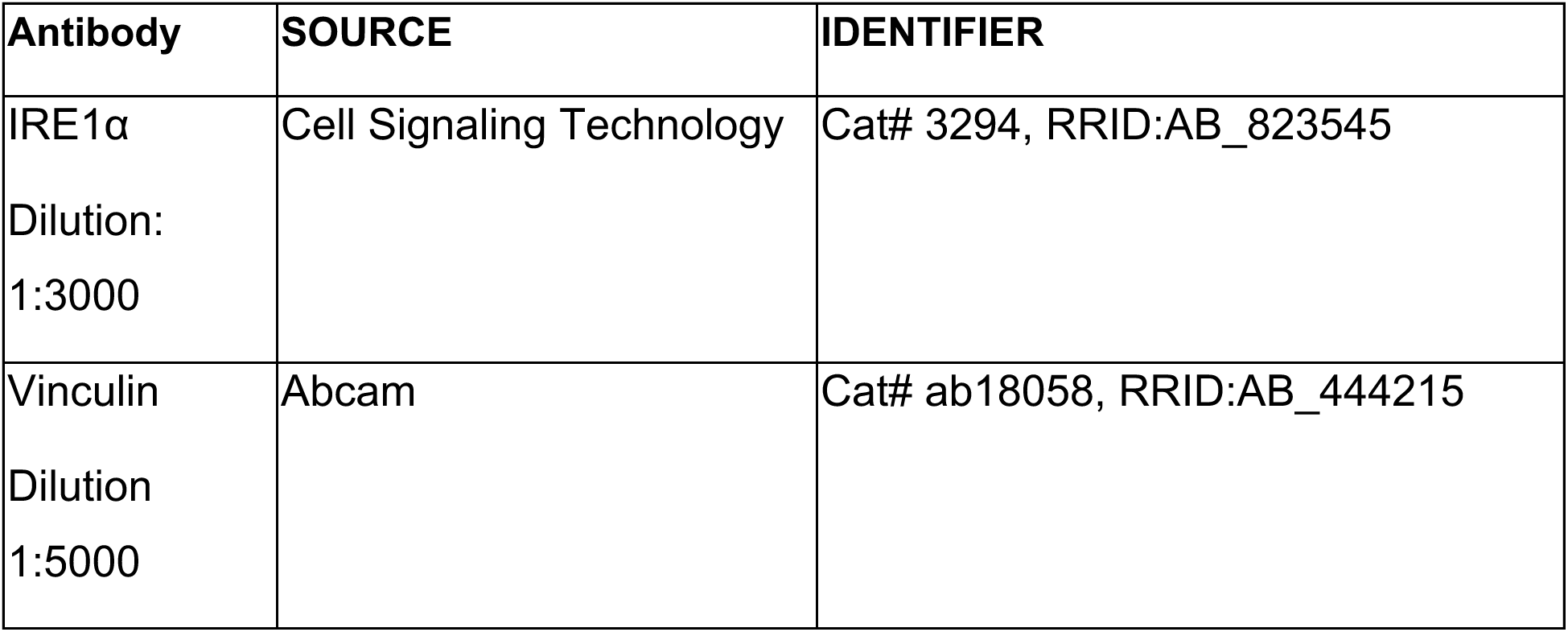
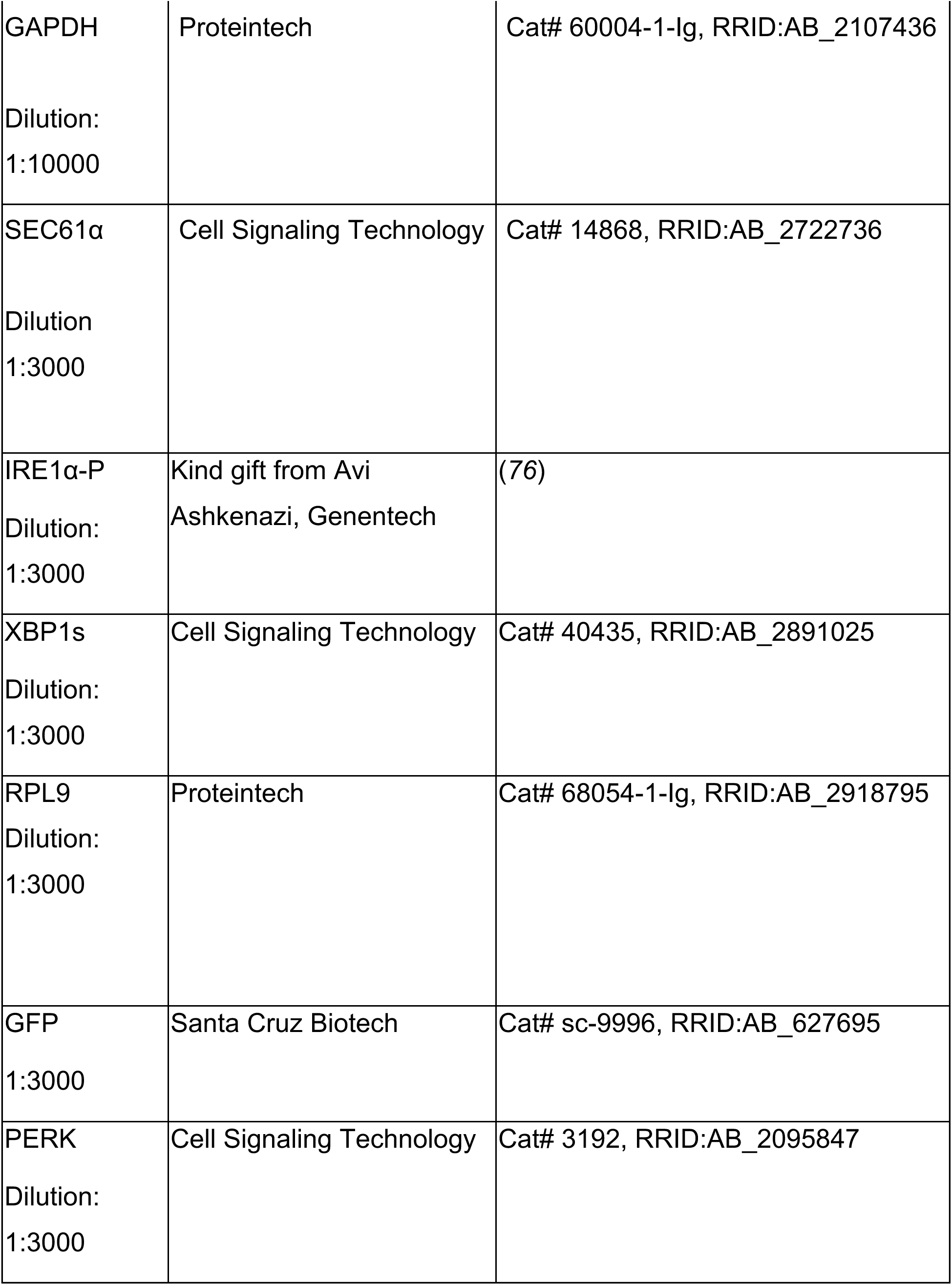
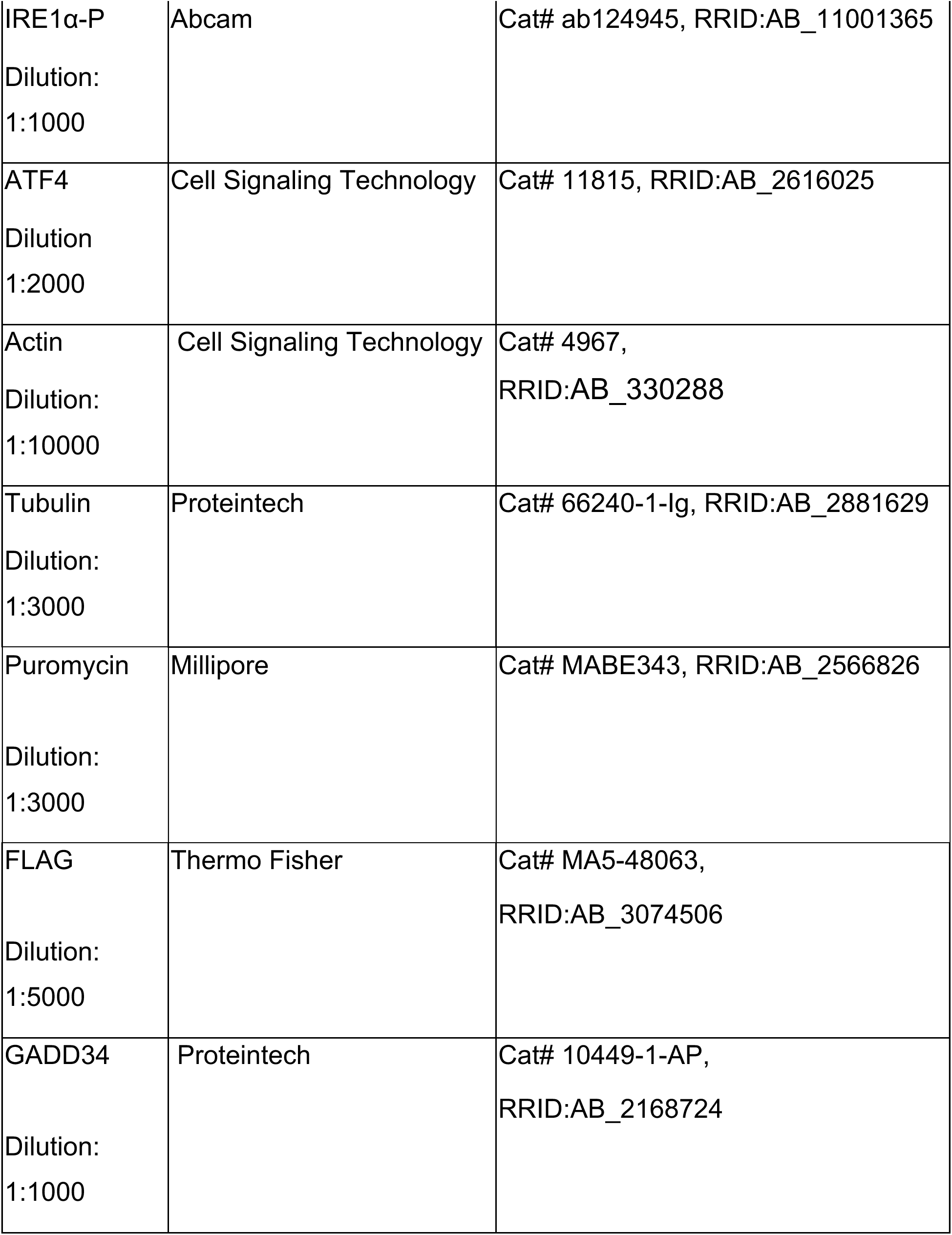
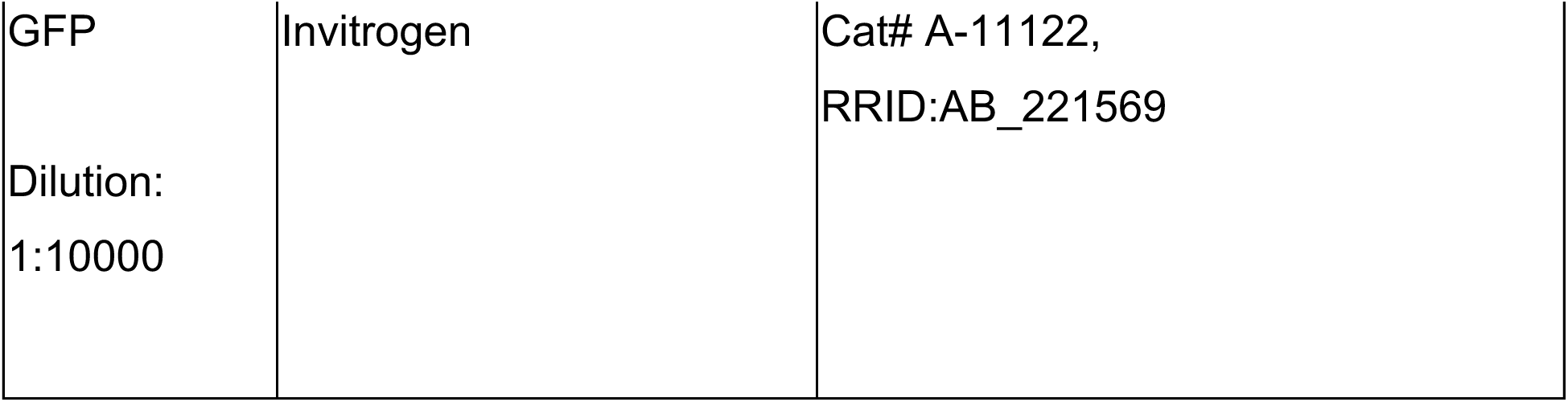
Antibodies used in this study.

### IRE1α immunoprecipitation

U-2 OS *IRE1α*^-/-^ cells expressing mNeonGreen-tagged IRE1α^WT^ or IRE1α mutants upon addition of 5 ng/mL doxycycline for 48 hours, or H4 cells expressing endogenous levels of IRE1α were washed three times in cold 1X PBS and lysed in a buffer containing 50 mM TRIS pH = 8, 150 mM NaCl, 1% digitonin, and protease inhibitor by end-to-end rotation for 30 minutes at 4 °C. The supernatant was collected after centrifugation of lysates at 15,000 rpm for 15 min.

For mNeonGreen-tag affinity mediated immunoprecipitation, 500 µg of clarified lysate was incubated with agarose or magnetic beads coupled to an mNeonGreen nanobody (ChromTek) overnight in a rotary shaker at 4 °C. Beads were then washed three times with a buffer containing 50 mM Tris pH 8, 150 mM NaCl, 0.1% digitonin, and protease inhibitor. Proteins bound to the beads were eluted by boiling the beads in elution buffer (20 µL of buffer containing 50 mM Tris pH 8, 150 mM NaCl, 0.1% digitonin, and protease inhibitor + 10 µL of Laemmli buffer + 2% 2-mercaptoethanol). The eluates were then subjected to SDS-PAGE and immunoblotting.

For endogenous IRE1α immunoprecipitation from H4 cells, lysates were first pre-cleaned with Protein A sepharose beads (Cytiva Cat no. 95016-992) on a rotating platform for 20 minutes at 4 °C. The samples were centrifuged at 5000 rpm for 5 minutes at 4 °C and the supernatant was collected. 500 µg of supernatant was then incubated overnight, rotating at 4 °C, with 2.5 µg of the anti-IRE1α antibody (Cell Signaling Technology). The beads were then washed 3 times with 800 µL of buffer containing 50 mM Tris pH 8, 150 mM NaCl, 0.2% digitonin, and protease inhibitor. The bound proteins were eluted from the beads by adding Laemmli buffer and boiling for 5 minutes at 95 °C. The eluates were then subjected to SDS-PAGE and immunoblotting.

### Live cell imaging

Time-lapse imaging was performed on U-2 OS *IRE1α-/-* cells expressing mNeonGreen-tagged IRE1α^WT^. Imaging was conducted using a Nikon CREST confocal spinning-disk system, equipped with a 40X water-immersion objective. Cells were maintained in a humidified chamber at 37 °C with 5% CO_2_ throughout the experiment. Images were acquired every 30 minutes for a total experimental period of 16 hours.

### RNA extraction, RT-PCR and quantification of *XBP1* mRNA splicing

To measure *XBP1* mRNA splicing in Figures 1D, 1G, S1B, S5A-B, 3C, 3E, 3G, 3K, S6B and S6J-K, total RNA was isolated using the RNeasy RNA purification kit (Qiagen) or the Promega Maxwell automated RNA extractor following the manufacturer’s recommendations. 1 µg of total RNA was reverse transcribed with SuperScript VILO (Invitrogen) following the manufacturer’s recommendations. The resulting cDNA was diluted 1:10 and used as a template and PCR was performed using primers listed in Table 3 and Hot Start Taq DNA polymerase (NEB). The PCR products were separated on 3% TAE-agarose gel.

**Table 3.**
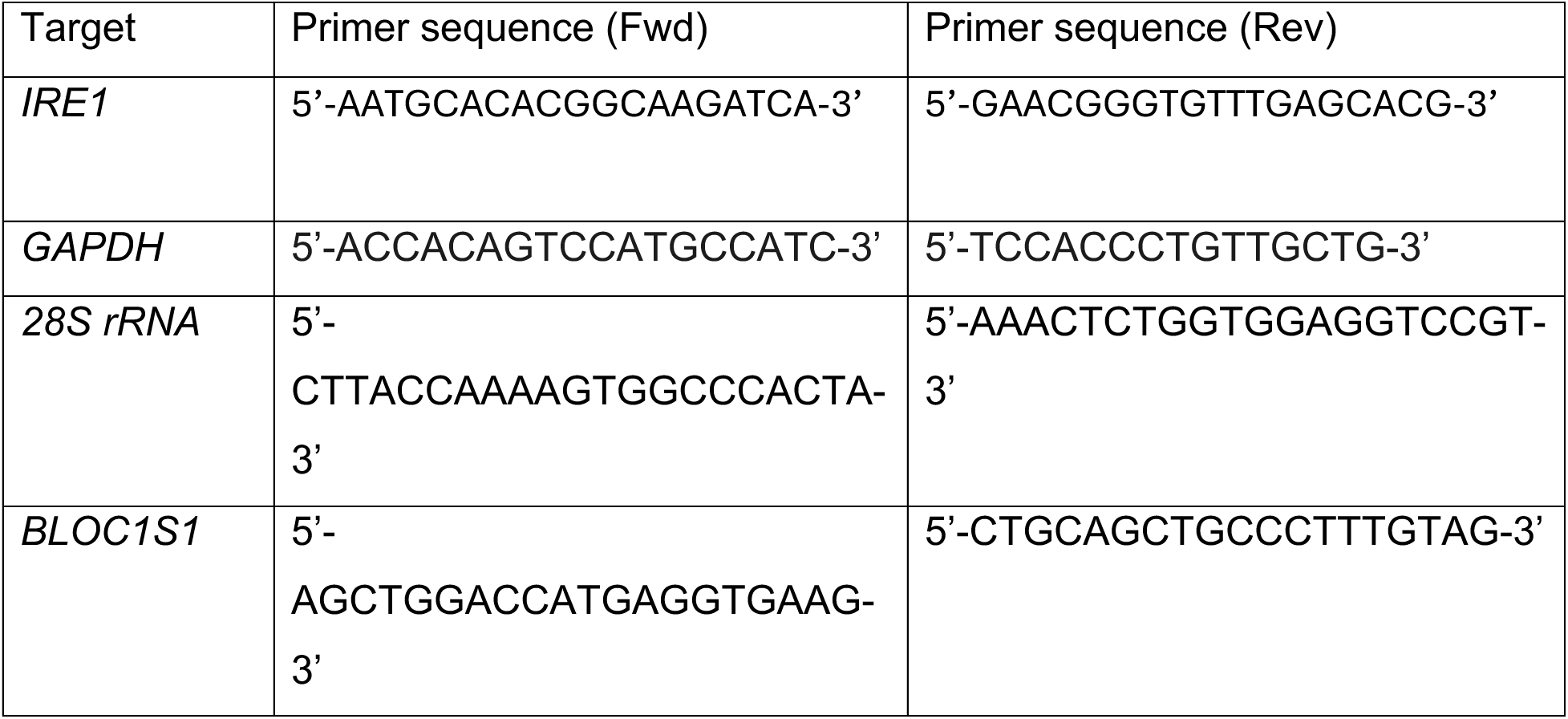
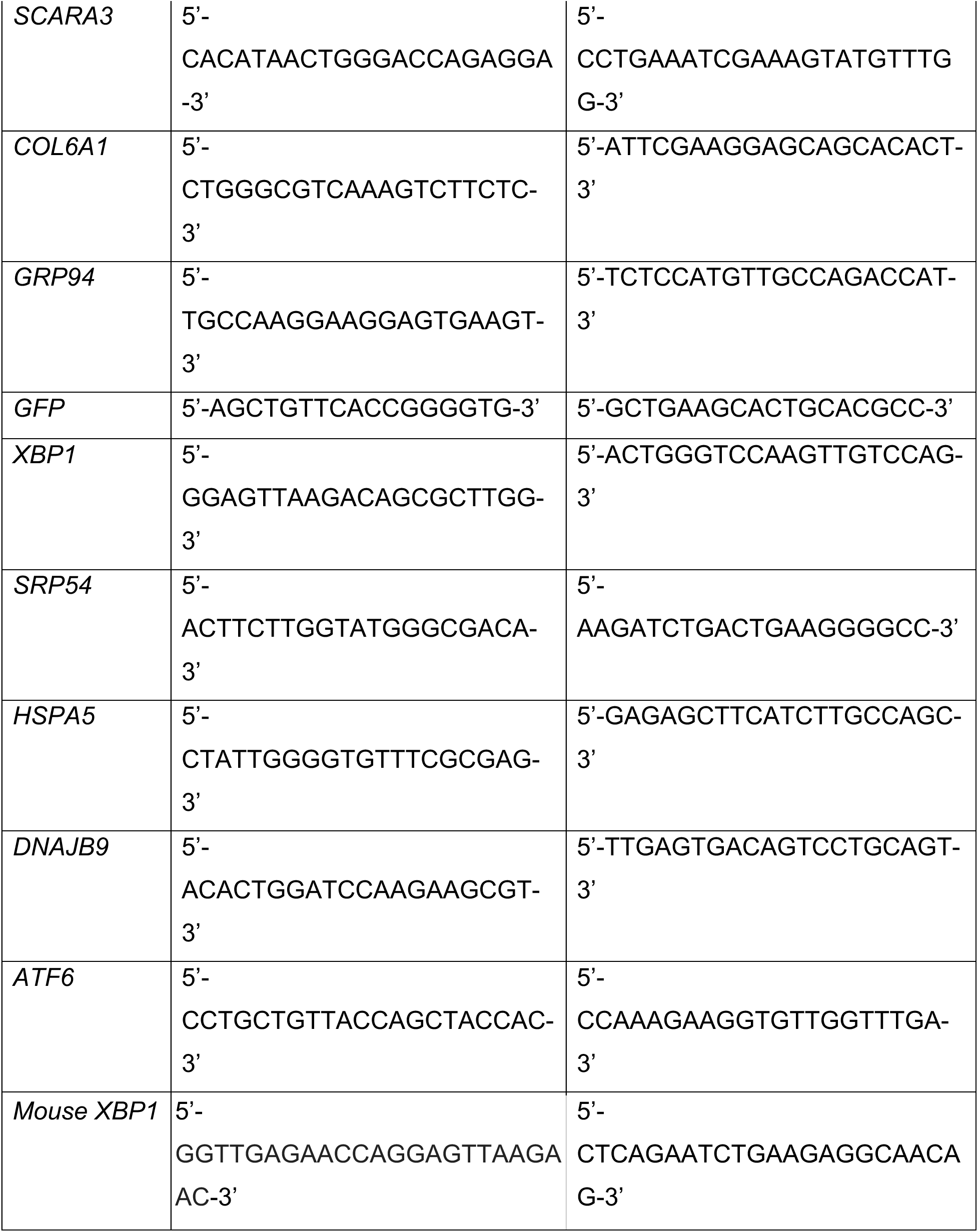
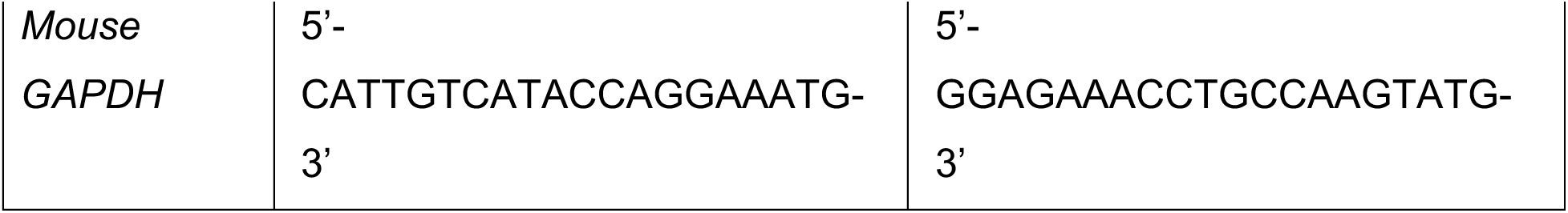
Primers used in this study.

To measure *XBP1* mRNA splicing in all other experiments, total RNA was isolated using the Direct-zol™ RNA Miniprep Plus kit (Zymo Research) according to the manufacturer’s protocols. The yield and the purity of RNA were determined by the spectrophotometer NanoDrop 2000. For RT-PCR analyses, 1 μg of RNA was converted to cDNA using the QuantiTect Reverse Transcription Kit (Qiagen). The cDNA was diluted 1:25 in the PCR reaction tube and the PCR was performed using Promega Go Taq (NEB). *XBP1* mRNA was amplified using the following primer pairs: *XBP1* Forward: 5’-TTACGAGAGAAAACTCATGGCC-3’; *XBP1* Reverse: 5’-GGGTCCAAGTTGTCCAGAATGC-3’. *GAPDH* mRNA was amplified using the following primer pairs: *GAPDH* Forward: 5’-ACCACAGTCCATGCCATC-3’; *GAPDH* Reverse: 5’-TCCACCCTGTTGCTG-3’. The PCR conditions used were as follows: Step 1: 94 °C - 3 minutes; Step 2: 94 °C - 30s; Step 3: 55 °C - 30s; Step 4: 72 °C - 30s; Step 5 - 72 °C - 7 minutes. Steps 2, 3 and 4 were repeated for 35 cycles. The amplified products were separated on a 3% TAE-agarose gel containing SYBR-Safe dye.

The gels were imaged using a BioRAD ChemiDoc gel apparatus system, and images were resized to enhance *XBP1*u and *XBP1*s band separation. For quantification, image analysis was performed using Fiji (ImageJ). Before quantification, background correction was applied, and a defined Region of Interest (ROI) was selected for each lane, encompassing all bands (spliced, unspliced, and hybrid - marked as * in the figure panels). The intensity was calculated for each band, and the percentage of spliced *XBP1* (*XBP1s*) was determined as follows: [(*XBP1*s + ½ *XBP1*h) / (*XBP1*s + *XBP1*u + *XBP1*h)] x 100. The *XBP1* hybrid band is a slower-migrating band that represents a hybrid DNA molecule. One strand of the hybrid is the unspliced (*XBP1*u) and the other is the spliced (*XBP1*s) form of the transcript, formed during the PCR reaction itself.

### RNA-seq and Bioinformatics Analyses

#### Library preparation

Libraries were prepared according to the Watchmaker mRNA Library Prep Kits (Watchmaker Genomics 7BK0001-096) protocol, using 200 ng of RNA input, and sequenced on a NextSeq 2000 using XLEAP-SBS chemistry.

#### Pre-processing and QC

Raw sequencing data were demultiplexed using bclconvert. RNA-seq FASTQ files were trimmed, aligned to the human reference genome (GRCh38, v108), and quantified using the nf-core/rnaseq pipeline with the star_salmon option (v3.12.0). Gene-level length-scaled counts were used for all downstream analyses.

#### Differential gene expression

Differentially expressed genes (DEGs) between conditions at each time point were identified within each dataset using the limma-voom framework. Raw length-scaled counts were filtered to retain genes with a minimum of 10 counts within biological groups and trimmed-mean of M-value (TMM) normalization factors were calculated. Empirical Bayes estimates of the prior variances were robustified against outlier genes. Thresholds of adjusted p-value < 0.05 after Benjamini-Hochberg correction and absolute log-fold-change > 0 were used to identify differentially expressed genes. Sample-wise quality weights were estimated and incorporated into the linear modeling for the FKBP-PKR dataset using the voomWithQualityWeights function.

#### Term Enrichment

Enriched Reactome terms within comparisons were identified using the continuous CAMERA test in the FKBP-PKR dataset and a hypergeometric test in the RocA dataset using the kegga function. The background for the hypergeometric test was all genes that were tested for differential expression. Significant enrichments were identified with an adjusted p-value < 0.05 after Benjamini-Hochberg correction. To find groups of significantly enriched terms, we constructed a graph of Jaccard similarities between all significant terms based on the number of constituent genes, removed edges with similarities below 0.3, and performed Louvain clustering with a resolution of 1.

#### XBP1 splicing

To quantify XBP1 slicing (*XBP1s*) levels, salmon was run with 100 bootstraps for each sample, and transcript-level overdispersion parameters associated with quantification uncertainty were estimated using the catchSalmon function in edgeR. Raw counts were corrected for overdispersion, transcripts with fewer than 10 counts across biological groups were removed, and TMM normalization factors were calculated. Counts-per-million (CPM) were calculated per sample, and slicing levels were measured as ratios of CPMs between XBP1s (ENST00000344347) and the sum of all annotated Ensembl XBP1 transcripts.

### RT-qPCR analyses

Total RNA from sub-confluent H4 cells or from U-2 OS cells was isolated using the RNeasy RNA purification kit (Qiagen) or the Promega Maxwell automated RNA extractor following the manufacturer’s recommendations. 1 µg of total RNA was reverse transcribed with SuperScript VILO (Invitrogen) following the manufacturer’s recommendations. The resulting cDNA was diluted 1:10 and used as a template for qRT-PCR using PowerUp SYBR Green Master Mix (Applied Biosystems) according to the manufacturer’s protocol. *GAPDH*, *ACTIN*, or 28S rRNA was used as a normalizing control to estimate fold changes in mRNA expression.

### Molecular docking of SEC61-IN-1 into the SEC61 translocon

The compatibility of SEC61-IN-1 (SEC61-IN-1/A317) to fit into the canonical SEC61 inhibitor site (*49*) was assessed using interactive docking by analogy to the existing compounds ipomoeassin F (PDB ID pdb_0008do1) and eeyarestatin I (pdb_00008do3). In brief, the protein component of the eeyarestatin I complex was prepared for simulation in ISOLDE (*77*) by deleting the existing ligand, adding hydrogens, and restraining the geometry with a web of local distance restraints using the “Isolde restrain distances” command. SEC61-IN-1 was parameterised for simulation using the GAFF2 force field and placed as a rigid body just outside the binding pocket. The ligand was then maneuvered into the pocket in an interactive physics-based simulation in implicit solvent and allowed to settle into position without further restraints. The resulting model buries approximately 460 Å^2^ without significant change to the protein coordinates. We note from existing structures that this site in SEC61 exhibits considerable conformational flexibility; it is likely that binding of this compound would involve some further rearrangement of the protein, but we have not attempted to model that here.

